# An approach to produce thousands of single-chain antibody variants on a SPR biosensor chip for measuring target binding kinetics and for deep characterization of antibody paratopes

**DOI:** 10.1101/2025.01.11.632576

**Authors:** Rebecca L. Cook, William Martelly, Chidozie V. Agu, Lydia R. Gushgari, Salvador Moreno, Sailaja Kesiraju, Mukilan Mohan, Bharath Takulapalli

## Abstract

Drug discovery continues to face a staggering 90% failure rate, with many setbacks occurring during late-stage clinical trials. To address this challenge, there is an increasing focus on developing and evaluating new technologies to enhance the “design” and “test” phases of antibody-based drugs (e.g., monoclonal antibodies, bispecifics, CAR-T therapies, ADCs) and biologics during early preclinical development, with the goal of identifying lead molecules with a higher likelihood of clinical success. Artificial intelligence (AI) is becoming an indispensable tool in this domain, both for improving molecules identified through traditional approaches and for the de novo design of novel therapeutics. However, critical bottlenecks persist in the “build” and “test” phases of AI-designed antibodies and protein binders, impeding early preclinical evaluation. While AI models can rapidly generate thousands to millions of putative drug designs, technological and cost limitations mean that only a few dozen candidates are typically produced and tested. Drug developers often face a tradeoff between ultra-high-throughput wet lab methods that provide binary yes/no binding data and biophysical methods that offer detailed characterization of a limited number of drug-target pairs. To address these bottlenecks, we previously reported the development of the Sensor-integrated Proteome On Chip (SPOC®) platform, which enables the production and capture-purification of 1,000 – 2,400 folded proteins directly onto a surface plasmon resonance (SPR) biosensor chip for measuring kinetic binding rates with picomolar affinity resolution. In this study, we extend the SPOC technology to the expression of single-chain antibodies (sc-antibodies), specifically scFv and VHH, and dual-chain Fab constructs. We demonstrate that these proteins are capture-purified at high levels on SPR biosensors and retain functionality as shown by the binding specificity to their respective target antigens, with affinities comparable to those reported in the literature. SPOC outputs comprehensive kinetic data including quantitative binding (R_max_), on-rate (*k*_a_), off-rate (*k*_d_), affinity (*K*_D_), and half-life (*t*_1/2_), for each of thousands of on-chip sc-antibodies. Additionally, we present a case study showcasing single amino acid mutational scan of the complementarity-determining regions (CDRs) of a HER2 VHH (nanobody) paratope. Using 92 unique mutated variants from four different amino acid substitutions, we pinpoint critical residues within the paratope that could further enhance binding affinity. This study serves as a demonstration of a novel high-throughput approach for biophysical screening of hundreds to thousands of single chain antibody sequences in a single assay, generating high affinity resolution kinetic data to support antibody discovery and AI-enabled pipelines.

## Introduction

Biopharmaceutical companies invest billions of dollars annually in developing new drugs to prevent and treat diseases, yet only about 10% of drug candidates that enter development pipelines and clinical trials ultimately receive regulatory approval^1,2^. The majority of failures are attributed to inadequate efficacy and/or safety, which are often linked to the on- and off-target binding properties of drug molecules. As a result, there is a renewed emphasis on designing new molecular entities (NMEs), particularly biologics, that exhibit high affinity, specificity, and selectivity to their intended targets in early preclinical phase, towards furthering the candidates with improved profiles along the development pipelines. Currently, deep characterization of binding kinetics is typically performed during the lead selection process, and safety assessments are often conducted during costly in vivo animal studies (Figure 1). To improve success rates, it is critical to assess kinetic binding and safety profiles earlier in the preclinical workflow—ideally in the design phase—while striving to maintain a broader diversity of lead candidates. Achieving this requires new or improved methods to accelerate the design, build, and test cycles, for iterative improvement and down selection of most optimal lead candidates. Such advancements could significantly enhance clinical success rates, saving biopharma hundreds of millions of dollars annually, and expediting the delivery of new and improved therapies to patients.

**Figure 1:**
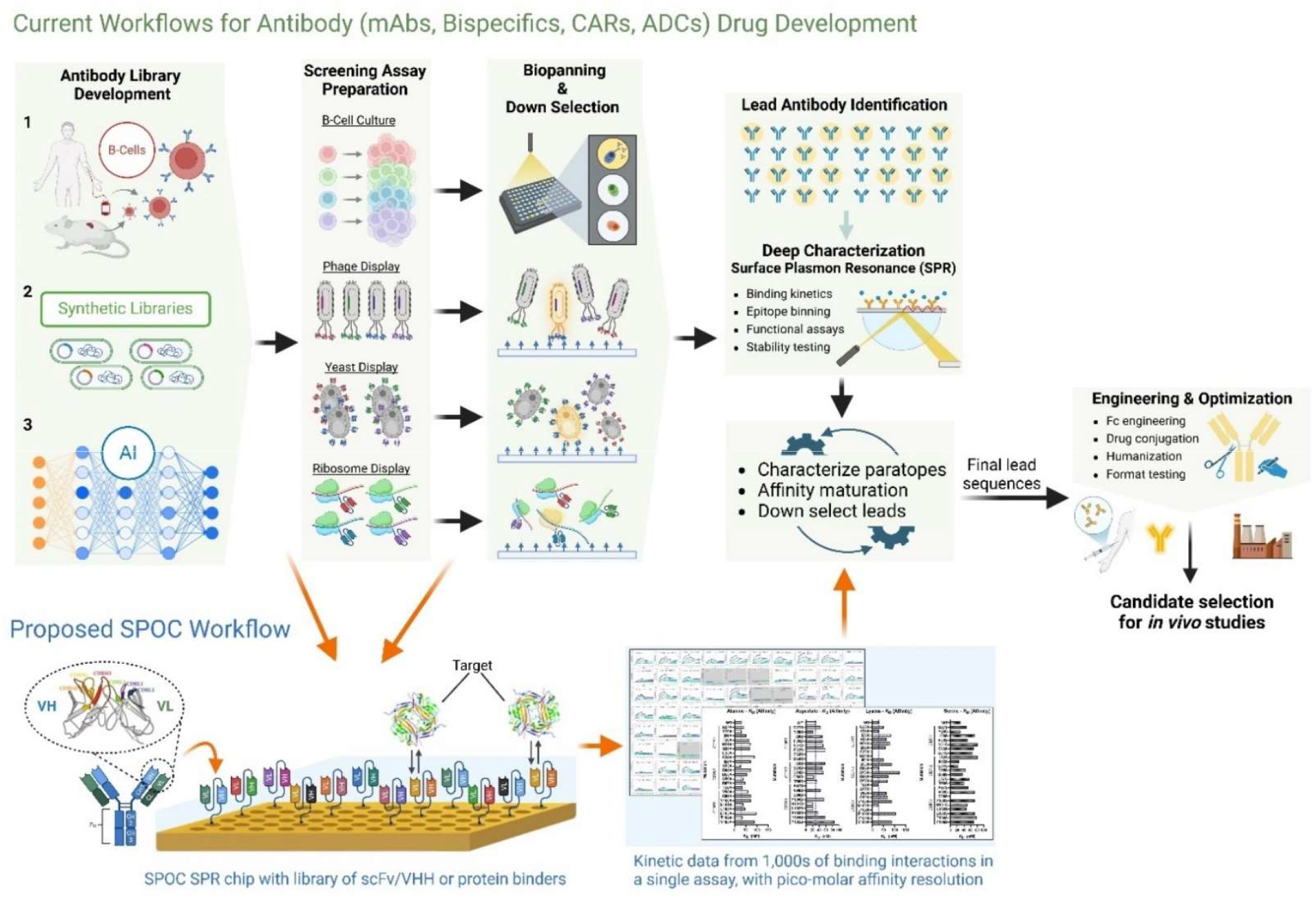
Schematic of current antibody drug development approaches and the proposed integration of SPOC into these workflows. Antibody sequence libraries have traditionally been harvested from 1) B-cells collected from naïve or immunized animals or convalescent patient serum, 2) derived from synthetic cDNA constructs, and 3) increasingly from generative artificial intelligence (AI) applied to in silico design of biologics. These inputs feed into a plurality of screening assays. Isolated B-cells are cultured, and secreted antibodies are assayed for clones that bind target antigen, which are subsequently sequenced. For inputs where sequences are already known, as after B-cell sequencing or for synthetic libraries or AI, various display technologies (phage, yeast, and ribosomes) can be directly employed whereby these cell-lines or ribosomes are applied to synthesize millions of sc-antibody variants for downstream panning against target antigens, and sequencing of the binders. After initial panning assays that isolate and identify the binders, deeper characterization is performed to assess the functional activity of lead candidates and interrogate their target binding mechanisms and kinetics with the use of label-free techniques such as surface plasmon resonance (SPR). These workflows involve additional paratope characterization steps and affinity maturation cycles, with the objective of down-selecting final lead sequences that are further engineered and optimized for efficacy and safety, prior to initiating humanization, Fc engineering and in vivo studies. We propose integrating SPOC platform into current workflows (lower part of the figure), wherein variable domain sequences derived from libraries can be directly leveraged for cell-free production and capture of sc-antibody libraries on SPR biosensor chips. SPOC facilitates measuring binding kinetics of all binders for down-selection by producing rich data, enabling deep characterization of the paratopes of lead antibodies and antibody affinity maturation cycles towards selecting optimally engineered lead candidates.

Machine learning (ML) and generative artificial intelligence (AI) are revolutionizing drug development, with several AI-designed drug molecules now advancing to clinical trials, signaling a transformative era in drug design and discovery. Current ML methods for drug discovery leverage iterative design-build-test-learn (DBTL) cycles to create high-affinity binders for drug targets (Figure 2). AI models can produce millions of initial candidate binders in a single ‘design-phase’ run, that are reduced to a few thousand by *in-silico* binding predictions. However, the subsequent build (synthesis) and test phases remain major bottlenecks due to their ex-silico nature, which is both capital and time-intensive. This capability-gap limits the scalability of DBTL cycles, as only a fraction of the candidates can be synthesized and tested experimentally. While powerful, AI models for protein design and binding analysis rely heavily on high-quality, large-scale wet lab data sets to train and refine their predictions, and are thus only as good as the real-world data they are trained on. New wet lab technologies are needed not only to generate data at scale for initial training but also for testing and improvement of antibody and protein binders through iterative cycles.

**Figure 2:**
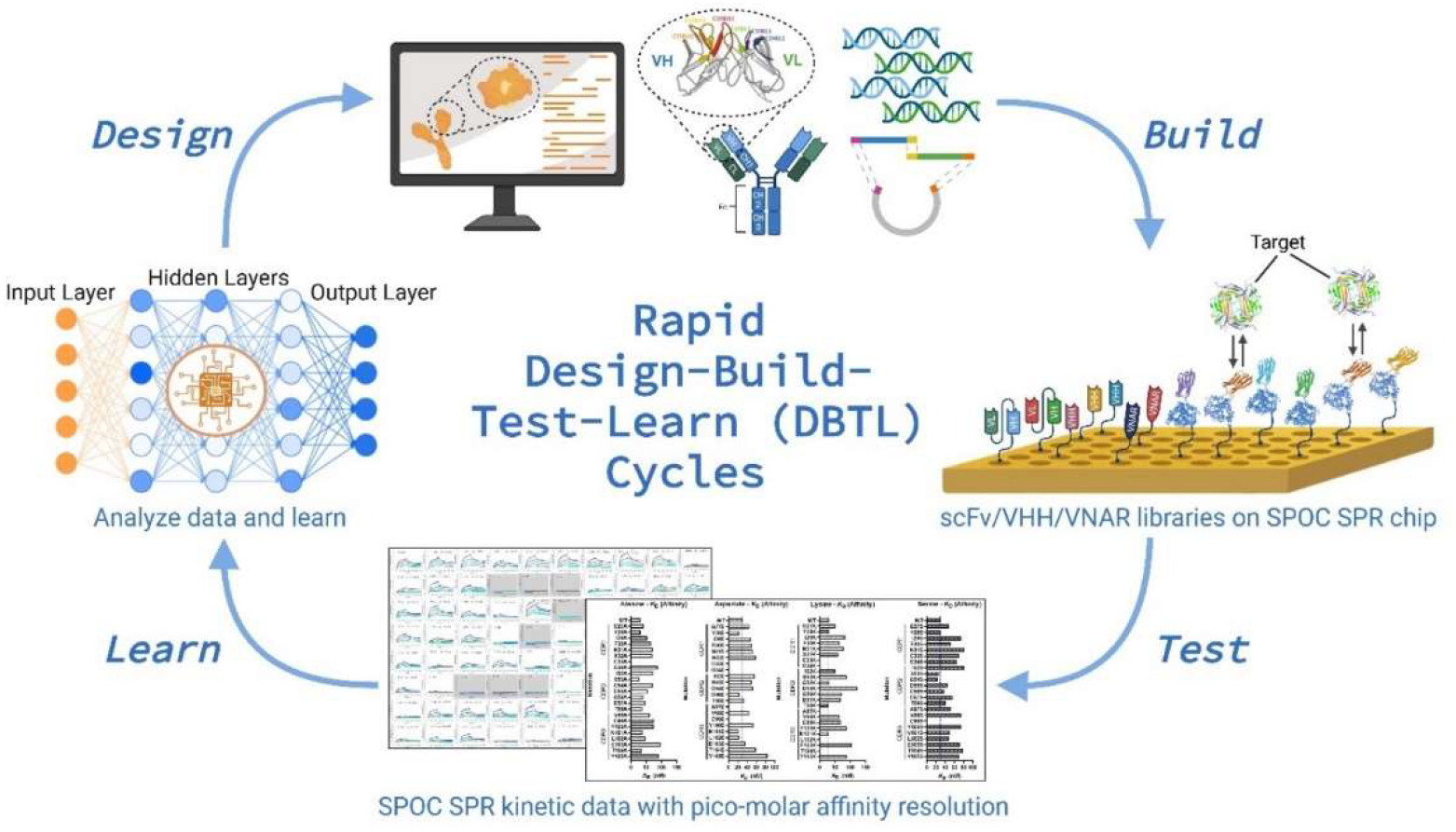
Schematic of SPOC assay integration into iterative AI-driven discovery workflows. Using inputs from trained AI models, sequences for numerous antibody sequences or protein binders (or other target-binding scaffolds and protein designs) can be cloned into cell-free expression vectors to rapidly build sc-antibody or protein libraries on SPOC SPR chips that can be screened with the desired target(s) to yield a wealth of detailed kinetic binding data. These rich kinetic datasets can then feed back into AI models providing additional training data to learn from, enabling the further refinement of AI-designed binders.

For conventional and AI-driven drug discovery workflows, experimental testing methods typically balance between ultra-high-throughput assays, which offer binary yes/no binding data, and more comprehensive techniques that yield detailed kinetic and biophysical data but can only accommodate a limited set of binders. Due to the cost-prohibitive nature of synthesizing and testing thousands of candidates, biophysical characterization is often limited to a few dozen lead molecules, representing a significant forced-reduction in sequence diversity. For example, interrogating binding characteristics using techniques such as surface plasmon resonance (SPR) in a high throughput manner is limited both by the limitations in the build (synthesis) phase as well as requirement for significant quantities of each drug candidate for sensor deposition and testing, limiting throughput. Moreover, ultra-high-throughput screening methods, such as phage and yeast display, often produce lead candidates with redundant sequences, reducing the diversity of final selections. The application of next-generation sequencing (NGS) has demonstrated its ability to identify rare clones missed by traditional pull-down assays, highlighting the inadequacy of current methods in capturing sequence diversity comprehensively^3–6^. These limitations underscore the need for novel high-throughput wet lab assays capable of producing diverse, high-quality data to better inform candidate selection and advance the integration of AI into drug development pipelines.

The class of antibody therapeutics and biologics has undergone significant innovation over the past two to three decades, driving the development of newer therapeutic modalities and yielding highly effective drugs, even as we enter the new age of AI-driven drug design. Following the success of full-length antibody therapeutics such as trastuzumab (Herceptin), bevacizumab (Avastin), and adalimumab (Humira) at the turn of the century, the therapeutic landscape has expanded dramatically to include a diverse and powerful array of novel Fab (dual-chained antibody fragments) and single-chain antibody derivatives. These include single-chain antibody formats such as single-chain variable fragments (scFvs), single variable domain of heavy-chain antibodies (VHH, also known as nanobodies, trademark of Ablynx N.V), and shark-derived variable new antigen receptors (VNARs); this is in addition to antibody mimetics such as designed ankyrin repeat proteins (DARPins), and a diverse class of AI designed protein binders. Single-chain antibodies, in particular, offer several advantages due to their smaller size, which often enhances tissue penetration and have fewer functional components outside of the antigen binding region compared to most mammalian antibodies, potentially lowering immunogenicity. These compact structures are also easier to produce using a range of expression systems, including yeast and bacterial platforms such as *E. coli*.

Nanobodies, which are derived from the antigen-binding region of specialized single-domain antibodies found in camelids (such as llamas, camels, and alpacas), hold immense promise for current and future clinical applications. These compact molecules, ranging from 12 to 15 kDa in size, consist of a single immunoglobulin domain that exhibits high-affinity binding (nM to pM range) even though it lacks a light chain. The small size and unique structure of nanobodies enable them to access cryptic antigens that conventional full-length antibodies cannot reach. Additionally, in certain modalities, their compact nature can facilitate cellular uptake into the cytoplasm, allowing them to target intracellular antigens^7^. Like the variable regions of IgG molecules, nanobodies possess three complementarity-determining regions (CDRs) that form the antigen-binding site (paratope). However, unlike IgGs, nanobodies are more thermostable, less prone to aggregation, and exhibit a shorter half-life—though the latter can be extended through conjugation with specific proteins or fusion with albumin-binding VHH. The clinical potential of nanobodies was underscored in 2018 with the FDA approval of Caplacizumab, the first nanobody-based drug for humans, used to treat acquired thrombotic thrombocytopenic purpura^8^. Caplacizumab demonstrated not only therapeutic efficacy but also low immunogenicity, highlighting its safety for clinical use. Since then, nanobody-based therapies have rapidly expanded with a CAR-T cell therapy (Ciltacabtagene autoleucel) incorporating a nanobody-based antigen receptor that was approved in the US and EU in 2022, a nanobody for treating solid tumors (Envafolimab) approved in China in 2021, and Ozoralizumab, a nanobody to treat rheumatoid arthritis approved in Japan in 2022^9–11^. Numerous other nanobody based therapies are in clinical trials for treatment of infectious disease, autoimmune disease, and cancer.

Given the therapeutic promise of sc-antibodies and protein binders, drug developers are increasingly focusing on engineering these molecules to address previously encountered challenges related to affinity, specificity, selectivity, and tissue distribution observed in earlier generations of antibody drugs. These efforts aim to enable access to previously obscure targets or specific epitopes, significantly expanding therapeutic possibilities. The growing popularity of sc-antibody drug discovery is evident from the proliferation of commercially available synthetic libraries, including fully naïve libraries, designed for rapid target screening and high throughput binder identification. AI-driven engineering of sc-antibodies and protein binders offers the potential to further accelerate the development of new therapies by optimizing paratopes for specific clinical targets. However, realizing this potential requires the development of new wet lab techniques capable of high-throughput, deep characterization of these engineered molecules. Such techniques are essential for rapid testing, analysis, and iterative improvement cycles to identify leads with very high affinity (or affinity optimized for specific therapeutic modalities), high specificity, and high selectivity, ensuring their readiness for preclinical and clinical testing.

As part of design-build-test-learn (DBTL) cycles in drug development, affinity maturation campaigns are frequently undertaken to iteratively enhance the binding properties of identified binders and lead candidates. The initial step in this process involves deep characterization of the full length and/or sc-antibody paratopes. This is typically achieved through comprehensive degenerate mutagenesis or computationally driven designer mutations introduced into the CDRs, linkers, and/or scaffold regions. Again, current limitations in the “build” and “test” phases significantly constrain this process. The high cost of producing the library of mutants designed by AI limits the number of sequences that can be experimentally tested for lead selection. As a result, mutations are often restricted to specific substitutions, such as alanine scanning. This narrow approach risks overlooking critical paratope residues essential for binding or alternative amino acid substitutions that cumulatively could significantly enhance binding affinity, potentially missing opportunities to achieve sub-picomolar binders and to develop “best-in-class” drug candidates. The limitations of current technologies in generating and testing sufficient sequence diversity can result in suboptimal lead candidates, undermining the potential of affinity maturation campaigns. To address these challenges and bridge the gaps in the “build” and “test” phases, we propose leveraging the previously reported Sensor-integrated Proteome On Chip (SPOC®) platform^12^.

The SPOC technology offers a transformative solution for high-throughput deep kinetic characterization of dual chain Fabs and sc-antibody variants (e.g., scFv and VHH), enabling the production of SPR biosensor chips with hundreds to thousands of unique antibody drug candidates produced and captured on chip. The platform allows for direct and simultaneous high-resolution kinetic measurements of analyte binding (e.g., antigen targets in solution) across the entire on-chip scFv, Fab, or VHH library in a single assay. To address the cost and scalability challenges of the “build” phase, the SPOC platform utilizes nanoliter-scale cell-free protein expression within high-density nanowells. This approach facilitates the production of thousands of properly folded proteins in discretely separated and isolated nanowells, within a 1.5-square-centimeter area, which are then directly capture-purified onto SPR biosensor chips. The SPOC system enables the characterization of binding interactions for up to 1000–2400 antibody variants against their antigen target or alternative analytes of choice (e.g., potential off-targets).

The sc-antibodies are covalently captured on the SPR biosensor chip, allowing for enhanced stability and offering the potential for multiple rounds of regeneration and follow-on assays. This feature ensures that a single chip may be reused for collecting replicate data, further improving cost-efficiency and throughput in the characterization and validation processes. Importantly, the SPOC workflow only requires the DNA sequence library encoding the sc-antibodies or protein binders, eliminating the need for expression and purification of proteins. By leveraging plasmid or linear DNA and cell-free expression systems to convert antibody gene/DNA libraries into protein libraries, the SPOC platform dramatically reduces the time and cost associated with obtaining high affinity resolution SPR kinetic data which provides a wealth of information about the binding characteristics of each sc-antibody on the sensor, including (R_max_), on-rate (*k*_a_), off-rate (*k*_d_), affinity (*K*_D_), and half-life (*t*_1/2_). Compared to traditional recombinant production approaches, this innovation accelerates the testing and validation process, making it feasible to generate the large-scale, high-quality wet lab data required to train AI models for better prediction accuracies. SPOC technology thus bridges critical gaps in the “build” and “test” phases, enabling rapid, cost-effective, and scalable characterization of antibody libraries to support state of the art drug discovery pipelines. In this study, we demonstrate the application of SPOC technology for the production and analysis of sc-antibodies and antibody fragments for the first time. The scFv, VHH, and Fab constructs tested and validated in this study are summarized in Figure 3. As a use case, we analyze the CDRs of a well-characterized HER2 nanobody utilizing single amino acid mutational scanning to identify key residues critical for optimizing binding affinity and function.

**Figure 3:**
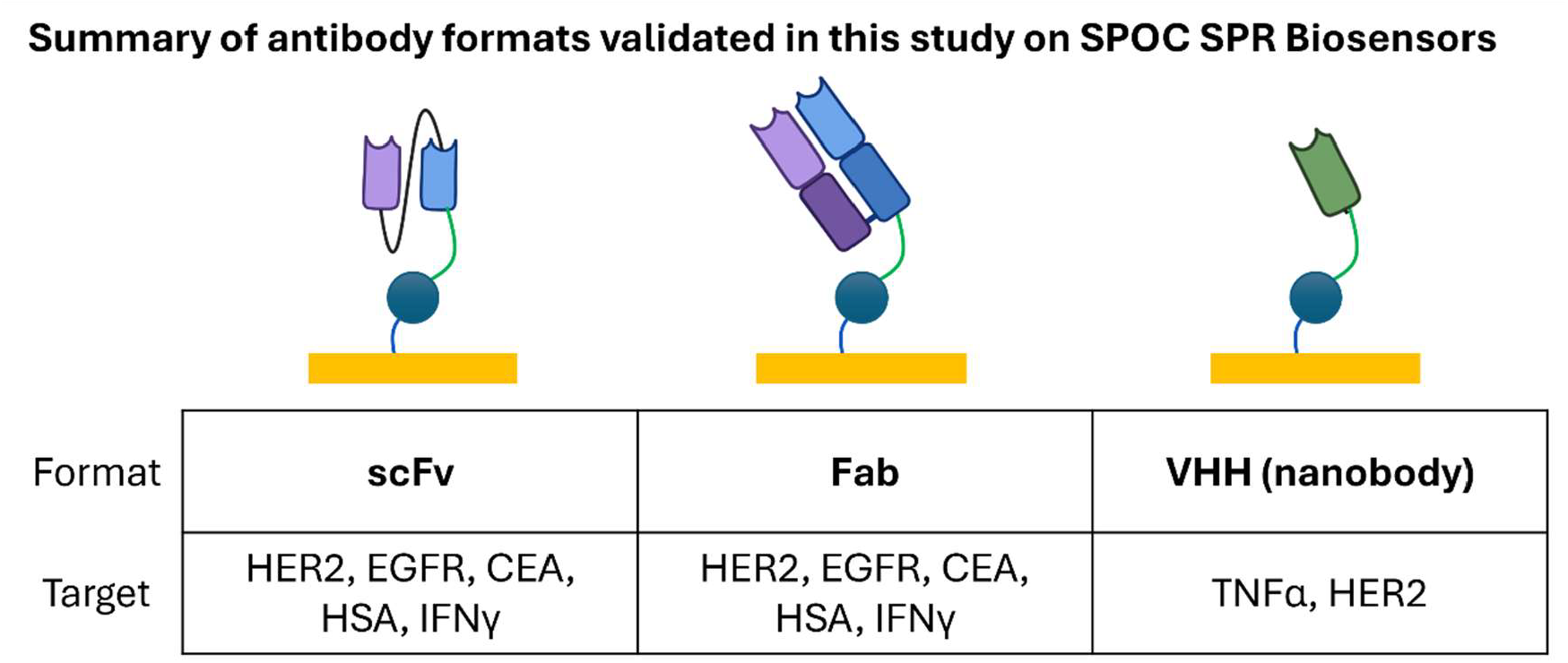
Summary of single chain and dual chain antibody formats validated in this study.

## Methods

### Materials and Reagents

Halo-PEG(2)-NH2*HCl (RL-3680) was sourced from Iris Biotech GmBH through Peptide Solutions, LLC. Rabbit anti-HaloTag (G9281) and TMR-Halo Ligand (G8251) were obtained from Promega. Mouse anti-HaloTag (#28a8) and Halo VHH (#ot) was obtained from ProteinTech.

Recombinant proteins for use as analytes and/or for epitope binning were sourced as follows: recombinant human TNF alpha (TNFα), BioLegend #570102; recombinant human IL-6, Biolegend #570802; recombinant human HER2 (AA 23-653), Acro Biosystems HE2-H5225; recombinant human CEACAM-5 (CEA), R&D Systems 4128-CM; recombinant human EGFR-Fc, R&D Systems 344-ER-050; recombinant human p53, Active Motif 8109; recombinant human interferon gamma (IFNg), BioLegend #570202; Trastuzumab, Selleck Chemicals A2007; Pertuzumab, Selleck Chemicals A2008; Rituximab, Selleck Chemicals A2009.

Detection antibodies were sourced as follows: Alexa Fluor® 647 Rat anti-human IL-6, Biolegend 501123; Mouse anti-human TNF-α Antibody, BioLegend 502901; Human anti-HER2 (Trastuzumab), Selleck Chemicals A2007; Human anti-CEA (Tusamitamab), Selleck Chemicals A2544; Mouse anti-p53, Sigma P6874; Goat anti-rabbit-Cy3 (Jackson ImmunoResearch 111-165-003), Goat anti-mouse-Cy3 (Jackson ImmunoResearch 115-165-062), Goat anti-Human IgG-Cy3, Jackson ImmunoResearch 109-165-098.

### Identification and construction of single-chain antibody sequences for testing

For proper evaluation of single chain antibodies on the SPOC platform, we chose to test sequences which were well characterized and reported in literature or were otherwise publicly available, such as FDA approved drugs. Sequences were identified using the ExPasy ABCD (AntiBodies Chemically Defined) Database (https://web.expasy.org/abcd/), the International Immunogenetics Information System (IMGT, https://www.imgt.org/), and associated PubMed resources.

Sequences were designed and codon optimized for cell-free expression using an IVTT *E. coli*-based kit. These constructs maintain a T7 Promoter-5’ UTR-Gene of Interest-HaloTag-T7 Terminator structure. The antibody sequences were derived from literature and converted to scFv format by extracting the VH and VL sequences and separating them with a (Gly_4_Ser)_3_ linker. A (Gly_4_Ser)_3_ linker was also placed between the scFv sequence and a C-terminal HaloTag. For the Fab format, distinct VH (fused to HaloTag) and VL DNA sequences were co-expressed in nanowells, that self-assembled and formed disulfide bonds, to form stable and functional Fabs. All DNA was synthesized by Twist Biosciences.

The HaloTag was chosen as the fusion tag of choice for our studies due to the covalent bond it forms with its chloroalkane ligand (Halo ligand) that is immobilized on the SPR biosensor surface. The HaloTag is derived from a halogenase enzyme; because the HaloTag protein contains an active site capable of catalyzing a covalent bond between the enzyme’s active site and a haloalkane when it is properly folded, the use of the HaloTag on the C-terminus of our constructs provides additional confidence that the linker-fused protein of interest is expressed in frame and is properly folded. The underlying rationale is that if the protein-Halo construct is covalently linked to a surface functionalized with chloroalkane (Halo ligand), the HaloTag itself must be folded properly to be enzymatically active for covalent bond formation with the chloroalkane on the sensor surface, and thus, the N-terminal fusion protein is more likely to have undergone proper folding as well.

### In-tube expression and gel validation of scFvs and nanobodies for validation studies

*E. coli* constructs were expressed according to the standard protocol for 6 hours at 37° C, and stored overnight at 4° C prior to assay the next day. For these constructs, a final DNA concentration of 10 nM was used and all reactions were performed in a 96-well PCR plate.

All HaloTagged products from *E. coli* expression reactions were visualized in-gel following fluorescent labeling of HaloTagged proteins via tetramethylrhodamine (TMR)-modified Halo ligand. Briefly, each cell-free expression reaction was incubated with TMR-Halo ligand in a 1:1:5 ratio of (undiluted expression reaction:50 nM TMR-Halo ligand (Promega):PBS) for 15 minutes at room temperature in the dark. Labeling with TMR-Halo ligand prior to denaturation for SDS-PAGE was critical to not destroy the Halo ligand active site/allow the covalent modification of the HaloTagged protein with TMR-Halo. Labeled samples were then prepared in both reducing and non-reducing conditions using Bolt LDS Sample Buffer (Invitrogen) and Bolt Sample Reducing Agent for reducing conditions (Invitrogen), incubated for 5 minutes at 95° C, and loaded into Bolt Bis-Tris 4-12% gels (Invitrogen). Gels were visualized using the fluorescent protein gel function on a Thermo iBright system (Thermo Fisher Scientific).

### Manual testing of single chain antibody binding

To analyze the ability of single chain antibodies to bind to their intended target and to measure any off-target binding prior to automated SPR analysis, scFvs and VHH were first tested manually in a multiplexed sandwich assay on glass slides. Glass slides with a hydrogel coating (Schott Nexterion Slide H) were first functionalized with 4 mM Halo ligand and then blocked with Superblock-TBS (Thermo Fisher Scientific). The slides were rinsed with diH2O and dried under a nitrogen stream, then affixed with a 64-well Flexwell incubation chamber (Grace Bio-Labs). 5 μL of HaloTagged scFv or VHH cell-free expression reaction was added to each well, followed by 15 μL PBST (PBS, pH 7.4, 0.2% Tween-20), and incubated for 1 h at room temperature on a rocker. Enough wells were filled to test each antigen against each scFv or VHH, regardless of specificity, to enable measurement of non-specific binding of antigen.

Wells were then washed 3 times with PBST, and the target antigen was added at a concentration of 50 nM for most, or 1:20,000 for IgG/IgM-depleted serum (Pel-Freez) for detection of human serum albumin. Negative controls were incubated with PBST instead of antigen. Antigen was incubated for 1 hour at room temperature on a rocker, then washed 3 times with PBST. Detection of antigen bound to scFvs or VHHs was done by incubating with a primary antibody against the antigen for 1 hour at room temperature on a rocker using a 1:500 dilution in 5% milk in PBST, then washed 3 times in PBST. For any primary antibodies that were not fluorescently labeled, a fluorescently labeled secondary antibody was incubated in the well for 30 minutes using a 1:500 dilution in 5% milk/PBST, then washed 3 times with PBST. One set of wells was also incubated with anti-HaloTag antibody and corresponding secondary antibody to measure total HaloTagged protein bound in the well. Prior to imaging, all slides were gently rinsed with diH2O and dried under a nitrogen stream.

Slides were imaged with an Innoscan 910AL (Innopsys) using sequential scanning of the 635 nm and 532 nm channels and Mapix software. Quantification was performed using a custom-made GAL file built for the 64-well Flexwell layout, in combination with the Mapix quantification feature. Background signal was subtracted from each well, then normalized to anti-HaloTag signal.

### Preparation and printing of nanowell slides

Silicon nanowell slides (1” x 3”) containing 11,280 individual nanowells were prepared as previously described for DNA printing^12^. Briefly, slides were treated with oxygen plasma for 2 minutes at 50 W followed by vapor phase coating with APTES ((3-Aminopropyl)-triethoxysilane) and curing for 1 hour at 100° C. DNA was diluted to 100 ng/μL in nuclease-free H2O. Printing of DNA into nanowells was performed with assistance from Engineering Arts, LLC using a Rainmaker 3 piezo-based printer (Bio-Dot) following printing of a print mix composed of bis(sulfosuccinimidyl)suberate (BS3, Thermo Fisher Scientific) and bovine serum albumin (BSA) into each individual well. Printed slides were allowed to dry in the humidified printing chamber for 15 minutes prior to removal, then stored in desiccating conditions at room temperature prior to assay.

### Capture of protein onto SPR biosensors

To enable capture of thousands of unique proteins onto planar substrates, SPOC Proteomics designed and built an in-house automated system termed Protein Nano Factory (PNF, previously referred to as AutoCap). The PNF system is capable of introducing cell-free expression lysate into each individual nanowell, incubating the slide for proper expression of protein within each nanowell, and transferring the expressed HaloTagged protein to a planar substrate (glass or biosensor) press-sealed against the nanowell slide^12^. The result is a planar substrate covered in discrete, circular spots each containing a pure protein uniquely expressed in each well, with no discernable cross binding between spots^12^. Xantec HC30M gold slides were used as the planar substrate for SPR biosensor production. Gold biosensor slides were first prepared for protein capture by activating for 10 minutes with a 1:1:1 solution of 0.4M N-(3-dimethylaminopropyl)-N-ethylcarbodiimide (EDC), 0.1 M N-hydroxysuccinimide (NHS), and 0.1 M 2-(N-morpholino) ethanesulfonic acid, pH 5.5 (MES), followed by water rinse and drying with a steady nitrogen stream. Slides were then functionalized with 1 mg/mL Halo-PEG(2)-NH2 overnight at room temperature (Iris Biotech; RL-3680). Any free NHS groups were quenched using 0.5 M ethanolamine, pH 8.5, then gently washed with diH2O and dried under nitrogen prior to loading onto the AutoCap instrument. Printed nanowell slides were prepared for the AutoCap instrument by blocking in SuperBlock-TBS for 30 minutes, then washed with diH2O and dried under nitrogen.

Following loading of the slides into individual chambers on the PNF system, the chambers were vacuumed and followed with injection of 500 μL of cell-free expression lysate. The nanowell and biosensor slides were then automatically pressed together following injection, isolating each individual nanowell with cell-free expression lysate and DNA to enable protein expression and immediate capture onto the biosensor. The chambers were incubated for 6 hours at 37° C prior to slide removal and immediate washing with PBST to remove unbound protein and lysate. SPR biosensors were either loaded immediately onto a custom Carterra LSA^XT^ SPR biosensing instrument for equilibration and assay, or stored in 50% glycerol at -20° C for later assay.

### Label-free and multiplex detection of in-solution analytes binding to single chain antibody molecules captured on SPR biosensors

Biosensors were rinsed with PBS followed by water prior to loading onto the Carterra SPR instrument compatible prism cartridge using 10-15 μL of refractive index-matching mounting oil (Cargille). The sensor was then loaded into a custom Carterra SPR instrument and equilibrated overnight in fresh, filtered, and degassed SPR buffer (1X PBS, 0.2% BSA, 0.05% Tween-20, pH 7.2). On this custom instrument, the individual protein spots are visible and can be assigned identifiers using regions of interest (ROI) prior to analyte screening. For validation of protein capture via mouse anti-HaloTag antibody binding, an association time of 6 min and dissociation time of 12 minutes was used. For analysis of analytes binding to single chain antibodies, 20 minutes association and 15 minutes dissociation was used. To calculate kinetic parameters for TNFα, a very high affinity nanobody, 1 hour association and 4 hours dissociation were used over two injections in regenerative conditions. For HER2 VHH wildtype and mutational analysis, 40 minutes association and 60 minutes dissociation were used to calculate kinetic constants using regenerative conditions.

For epitope binning experiments with commercial therapeutic antibodies, the screen was run in classical binning mode. The experiment cycles consisted of 30 min HER2 ECD antigen (20nM) injection followed by 15 min sandwich/binning therapeutic antibody injection (Trastuzumab [100nM], Pertuzumab [79 nM], or Rituximab [25 nM]) and a 15 min dissociation period. Two, 30-second pulses of 10mM Glycine-HCl (pH=3.0) regeneration buffer injections re-set the cycle after each dissociation phase between each therapeutic antibody tested. For epitope binning using VHH and scFv analytes expressed in crude IVTT lysate, the screen was run in standard kinetic injection mode, where a brief dissociation phase occurs between antigen and sandwich/binning analyte injections (in this case analyte being dilute, buffer exchanged IVTT lysate). Crude, 30μL IVTT reactions expressing HaloTag anti-HER2 VHH or HaloTag Trastuzumab scFv were first buffer exchanged into 1xPBS using Amicon 10k Ultra-0.5 Centrifugal Devices (89877) according to manufacturer instructions. IVTT binning screens were set to have 30 min association and 5 minute dissociation phases, with HER2 ECD antigen injected first at 20nM concentration followed by buffer exchanged IVTT lysate injection diluted 1:300 in running buffer. Regeneration (10mM Glycine-HCl, pH=3.0) was performed after IVTT lysate as a standard analyte injection before re-starting the cycle with subsequent rounds of antigen and IVTT lysate injections.

### Analysis of SPR biosensor data

Data collected from the custom Carterra LSA^XT^ SPR biosensing instrument was analyzed using Kinetics analysis software (Carterra). Raw data was y-aligned and double referenced against the leading running buffer only blank injection and no ligand control spots present on the sensor. Excessive spikes were filtered using standard settings (height = 7, width = 9). Processed, double-referenced data was then globally fit with a 1:1 Langmuir binding model to extract kinetic binding rates and other associated metrics [on-rate (*k*_a_), off-rate (*k*_d_), affinity (*K*_D_), R_max_, and half-life (*t*_1/2_)]. Mutants with approximately 10% of signal compared to the maximum RU signal observed for the highest concentration injection of 110 nM HER2 ECD were manually reviewed for consideration as a non-binder for purposes of analysis. Epitope binning experiments with commercially sourced therapeutic antibodies were analyzed using Epitope analysis software (Carterra) with only single-referencing to control ligand spots (no blank referencing) according to manufacturer instructions. Due to larger impact of bulk effect on signal for binning experiments performed with diluted, buffer exchanged IVTT lysate, double referencing was performed and the dataset was analyzed in Kinetics. Visualization of mutants was performed using PyMol version 3.1.3 based on PDB structure 5MY6^13^.

## Results

### Expression validation of scFv and VHH constructs

All scFvs and VHHs (Table 1) were designed as C-terminal HaloTag fusion proteins for expression using an *E. coli*-based IVTT expression kit lysate. All constructs were expressed via IVTT in 96-well PCR plate format and analyzed via SDS-PAGE in reducing and non-reducing conditions to confirm expression and molecular weight, and to evaluate cysteine bond formation. TMR-Halo ligand-labeled expression products were easily detected for all constructs at the expected molecular weight (Figure 4).

**Table 1:** List of scFv and VHH constructs tested

| Single Chain Antibody | References |
| --- | --- |
| Anti-Interleukin-6 (IL-6) (sirukumab) scFv | 14,15 |
| Anti-Human Serum Albumin (HSA) scFv & Fab | 16 |
| Anti-Tumor Necrosis Factor alpha (TNFa) VHH | 17 |
| Anti-Human Epidermal Growth Factor Receptor 2 (HER2) VHH | 13,18 |
| Anti-p53 scFv | 19 |
| Anti-Carcinoembryonic Antigen (CEA) scFv & Fab | 20 |
| Anti-EGFR (panitumumab) scFv & Fab | 21 |
| Anti-IFNγ (emapalumab) scFv & Fab | 22 |
| Anti-Her2 (trastuzumab) scFv & Fab | 23 |

**Figure 4:**
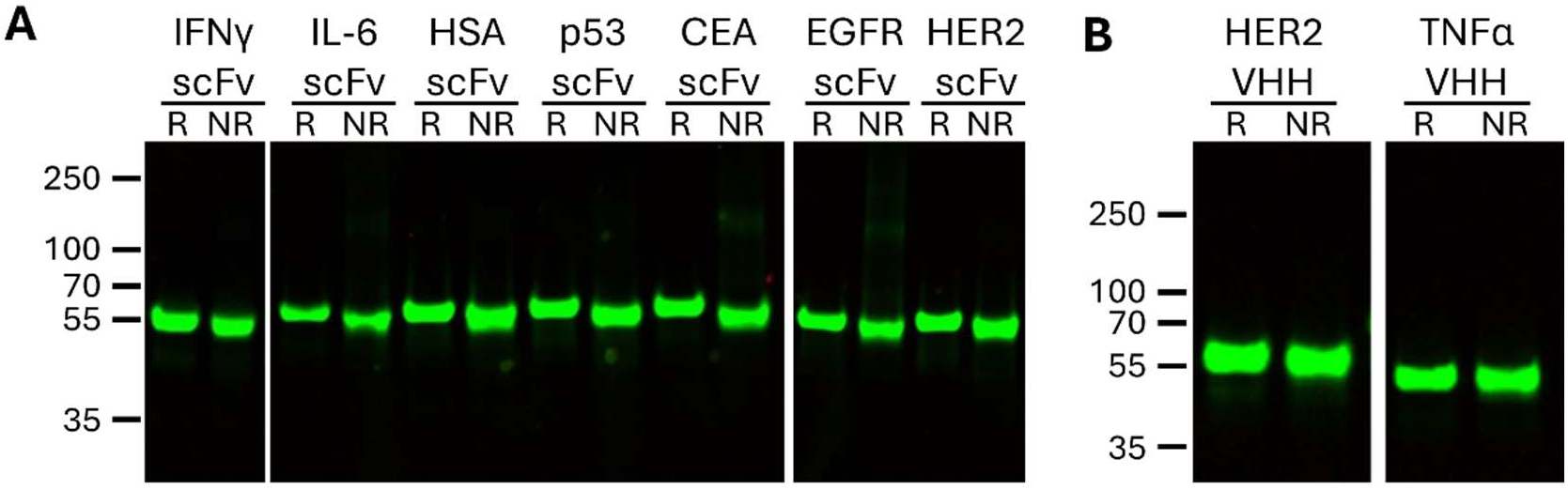
SDS-PAGE analysis of C-Terminal HaloTagged scFvs and VHH expressed using *E. coli* lysate. All expressed constructs were first covalently labeled with fluorescent Halo ligand then analyzed in reducing (R) and non-reducing (NR) conditions.

### Confirmation of antigen target binding

ScFv and VHH HaloTag fusions were assessed for binding and specificity for their target antigens in a fluorescent sandwich assay. Antigens bound their corresponding single chain antibodies at various levels but each scFv or VHH showed high specificity for its target compared to all other antigens tested (Figure 5). As expected, little to no signal was observed for the scFv or VHH constructs when antigen was omitted as a negative control (Figure 5A), nor when the incorrect antigen and corresponding detection antibody was used (Figure 5B), confirming specificity.

**Figure 5:**
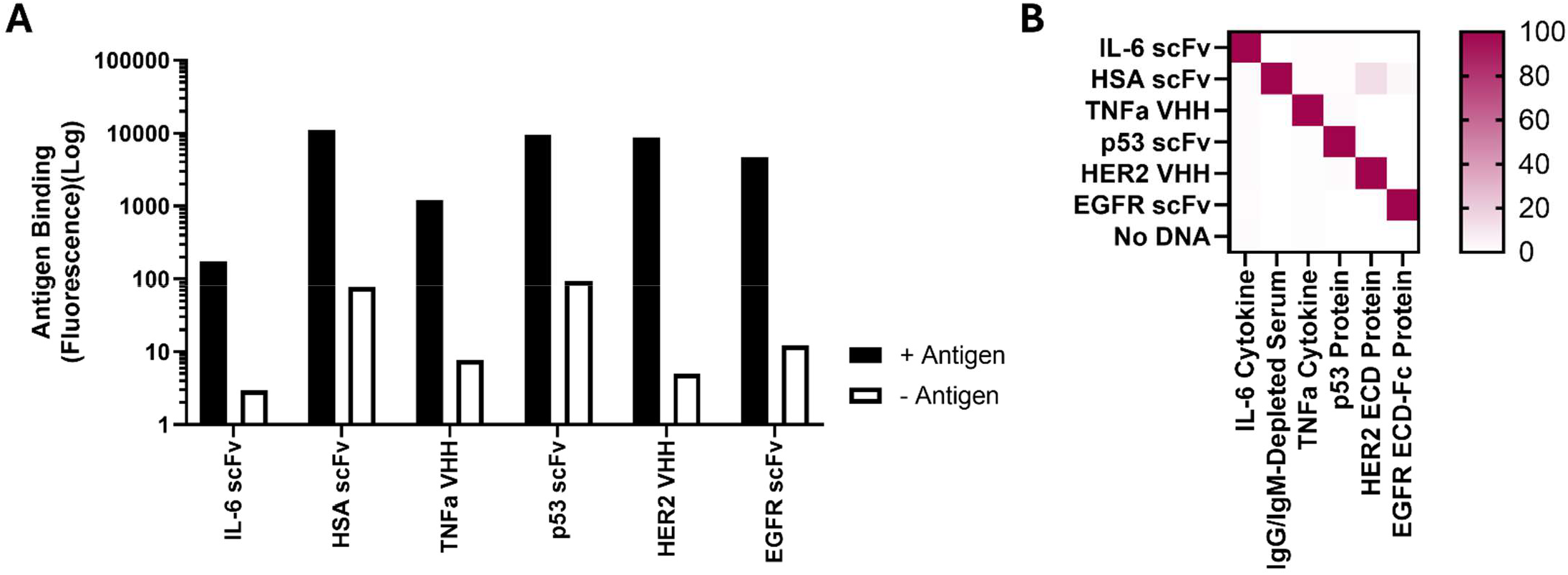
Fluorescent-based analysis of single chain antibody constructs binding to target antigen. C-terminal HaloTagged constructs were covalently linked to a glass surface via Halo ligand and incubated with (A) their target antigen (+Antigen) or PBST (-Antigen), or (B) all constructs were individually incubated with all antigens used in the assay. Antigens were detected via fluorescence using primary and secondary (if applicable) antibodies. IgG/IgM-Depleted Serum was used as the analyte for HSA.

### Expression, capture, and testing of single chain antibodies on multiplexed SPOC SPR biosensors

High density protein biosensors were prepared for evaluation of sc-antibodies expressed in nanowell slides printed with DNA encoding sc-antibodies constructs. ScFvs and VHHs expressed using an *E. coli* IVTT kit resulted in high levels (>200 RU) of protein captured on the surface, as measured with anti-HaloTag antibody (Figure 6).

**Figure 6:**
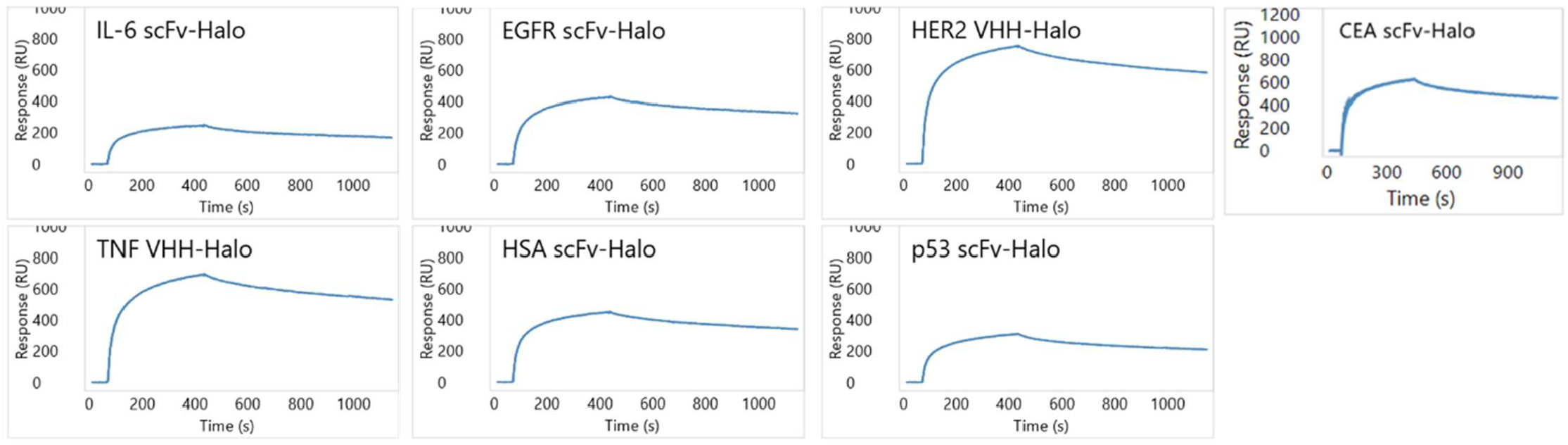
Confirmation of capture of scFv and VHH proteins with C-terminal HaloTag on SPOC biosensor. Capture of single chain antibodies on the biosensor following expression via *E. coli* IVTT lysate was validated via anti-HaloTag antibody via SPR.

Recombinant protein antigens matching each of the tested scFv or VHH constructs were flowed over the SPR sensor surface and binding levels and kinetics were measured. Multiple single chain antibodies, both in scFv and VHH formats (anti-CEA scFv, anti-EGFR scFv, anti-IFNγ scFv, anti-HER2 scFv, anti-HSA scFv, HER2 VHH, and TNFα VHH), were found to bind their target antigen at very high levels in a label-free format (Figure 7).

**Figure 7:**
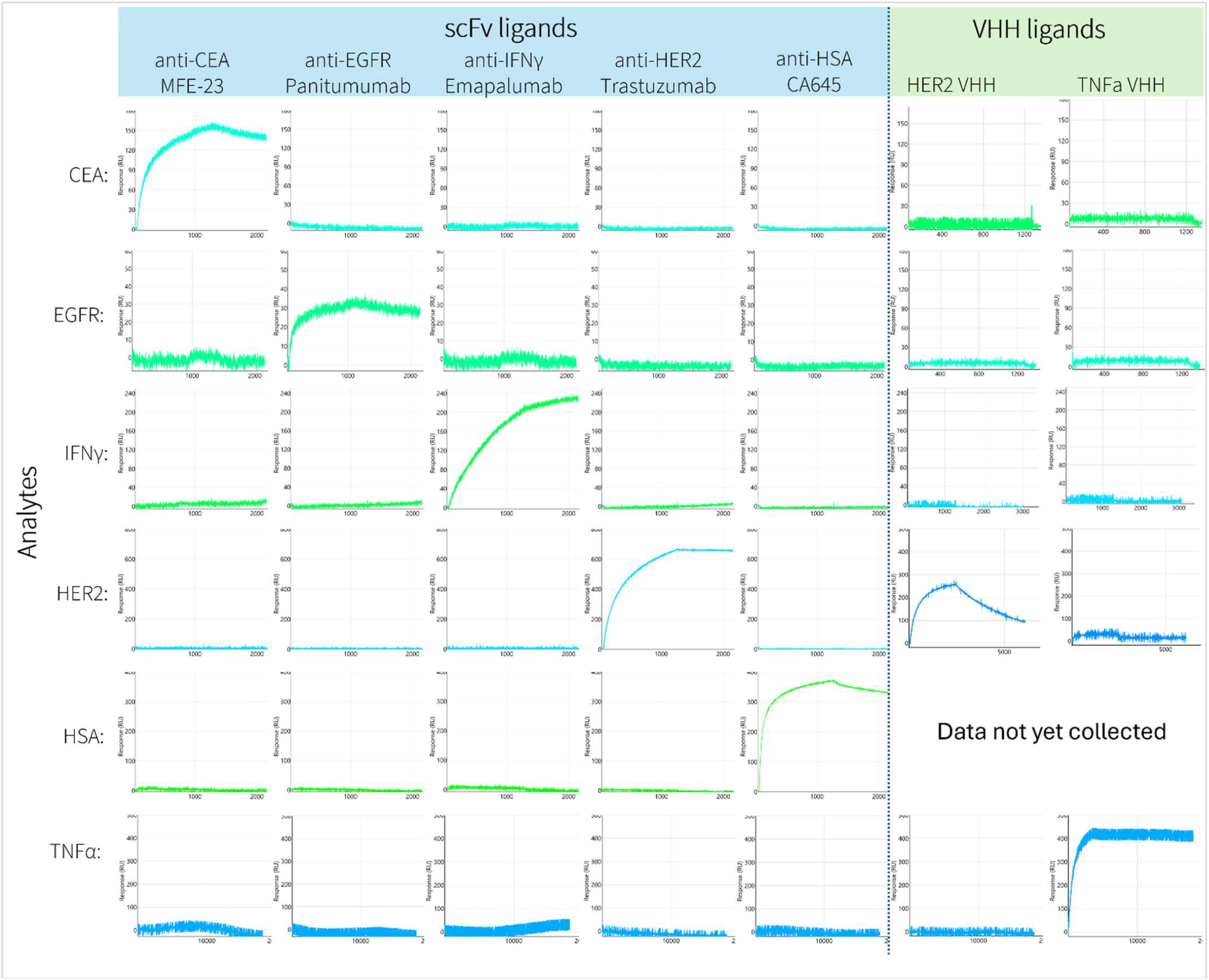
Analysis of scFv and VHH proteins on SPOC biosensor binding to antigen targets in comparison to non-targets. Antigen targets of the scFv and VHH proteins were analyzed for specific binding against all scFv and VHH simultaneously using a SPOC SPR biosensor. Binding traces to each cognate single chain antibody are shown. Data for scFv and VHH proteins were collected in two different SPR runs.

In addition to sc-antibodies, Fabs were also designed based on the same VH/VL pair used for scFvs in this study. Two separate DNA constructs encoding VH-CH1-HaloTag and VL-CL for each Fab were co-printed into nanowells and tested for co-expression/binding of antigen via SPR on SPOC biosensors. Preliminary testing indicated that antigens were specifically captured by the corresponding sensor-bound Fab (Figure 8).

**Figure 8:**
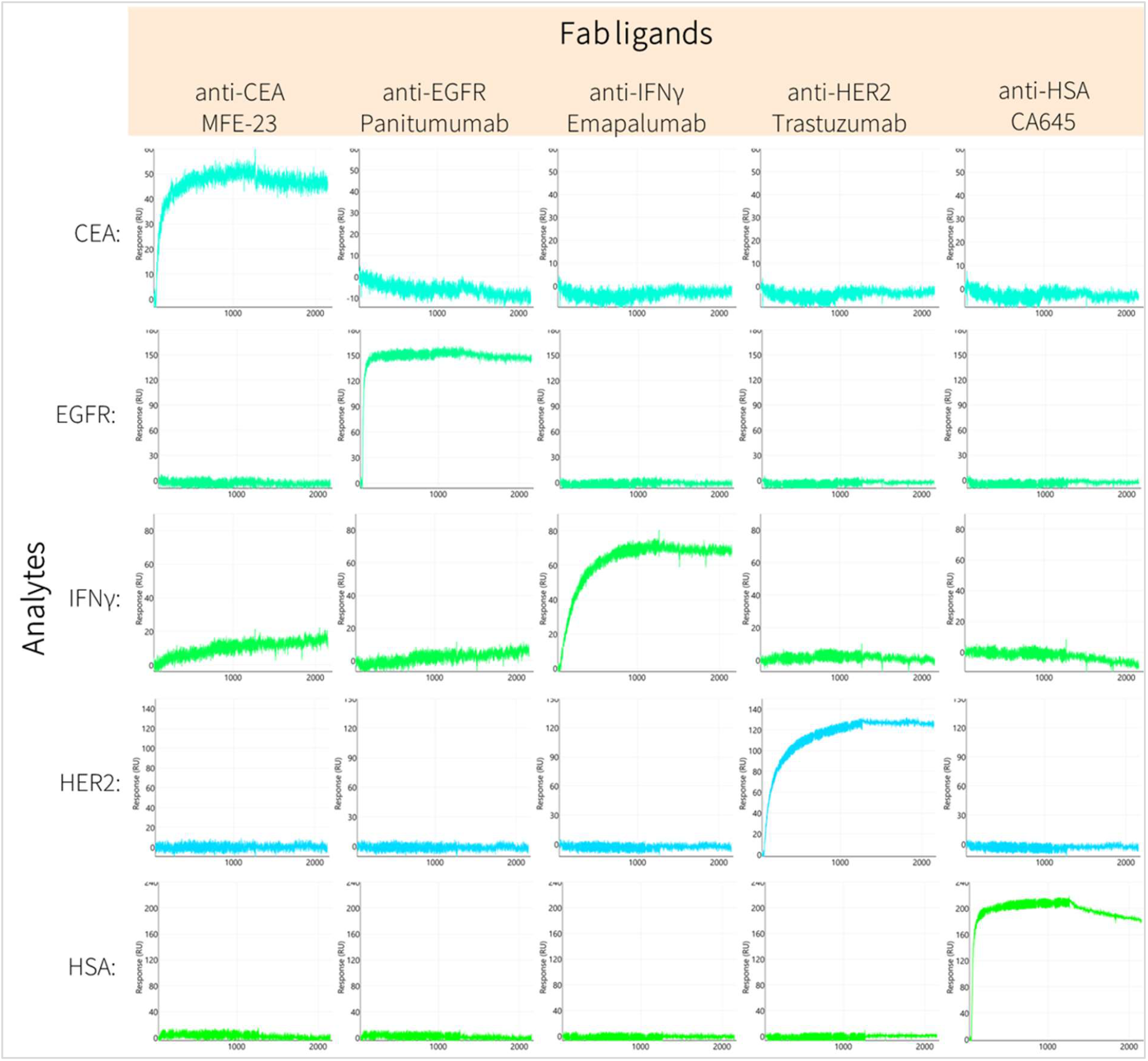
Analysis of Fab fragments on a SPOC biosensor binding to antigen targets in comparison to non-targets. Antigen targets of the Fab fragments were analyzed for specific binding against all Fabs simultaneously using a SPOC SPR biosensor. Binding traces to each cognate Fab are shown.

### Affinity calculations from TNFα VHH, HER2 VHH, and Trastuzumab scFv

Three constructs (TNFα VHH, HER2 VHH, and Trastuzumab scFv) were chosen for full kinetic analysis. Kinetics were measured on SPOC SPR biosensors in duplicate (replicate 1 and 2) by flowing recombinant antigen at multiple concentrations over the sensor surface while collecting kinetic data. Injecting TNFα over the biosensor using two concentrations (1 nM and 10 nM) and 4 hour dissociation, with regeneration, resulted in a calculated affinity (*K*_D_) of 149 pM (Figure 9A-B). This is 3-4 fold higher than the previously reported affinity of 540 pM for this VHH^17^. The affinity constant for HER2 binding to the HER2 VHH (Figure 9C-D) and HER2 scFv (Trastuzumab) (Figure 9E-F)was calculated to be 13.5 nM and 6.7 nM, respectively, using an increasing 5-step titration of HER2 antigen from 1.4 nM to 110 nM (Figure 9B). This compares well with the reported *K*_D_ of 4 nM in literature for HER2 VHH and 5 nM in FDA documentation for Trastuzumab.

**Figure 9:**
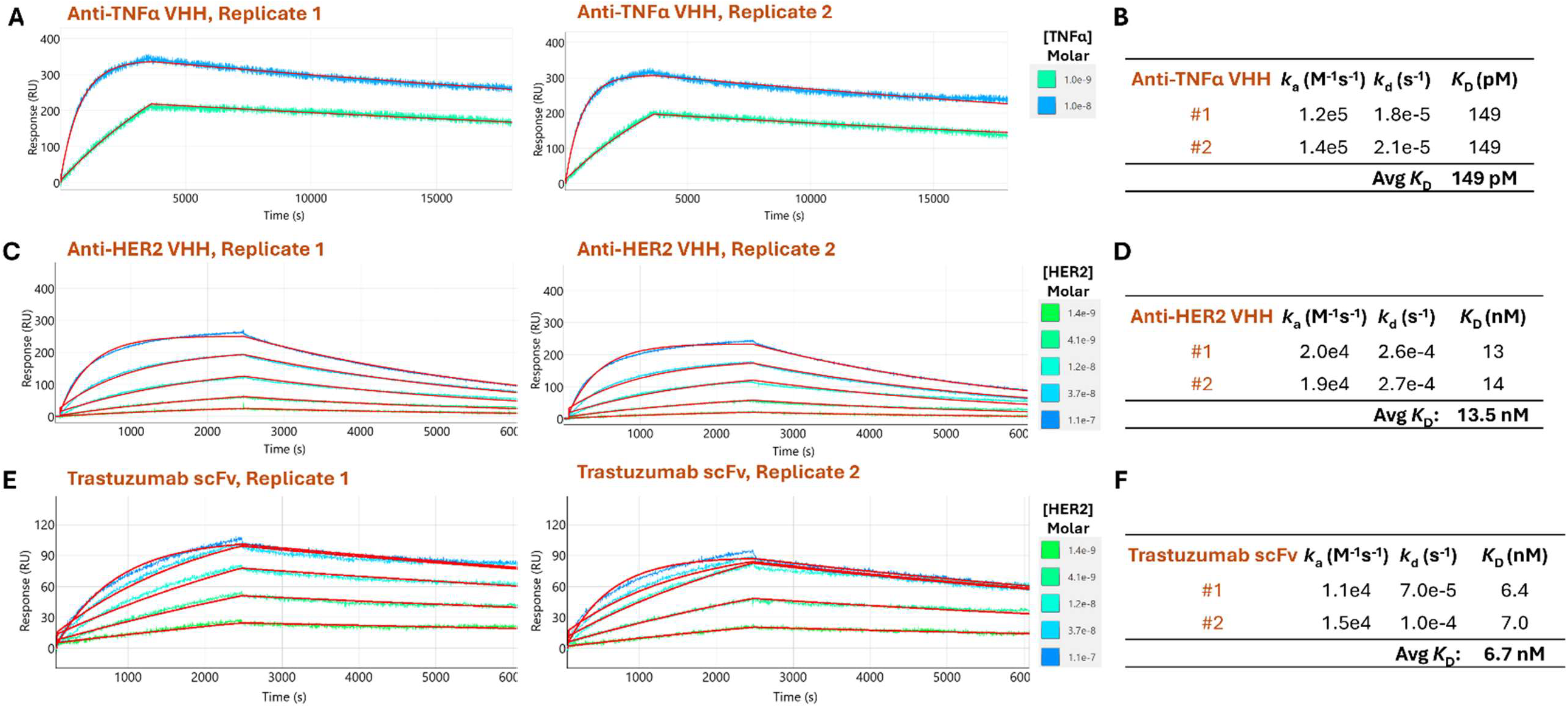
Kinetic analysis of TNFα and HER2 VHH constructs via SPR. Affinity constants for interactions between anti-TNFα VHH and TNFα were calculated from a titration of TNFα binding to the SPOC SPR sensor (A-B). Regenerative conditions paired with 4 hour dissociation was used between TNFα injections due to high affinity of the interaction. The affinity constant for the interaction between anti-HER2 VHH (C-D) and anti-Her2 scFv (Trastuzumab) (E-F) with recombinant HER2 was calculated from a titration of HER2 extracellular domain (ECD) binding using regenerative conditions. Kinetic measurements were averaged over the duplicate spots (1 and 2) for all three constructs.

### Epitope binning using purified sc-antibodies and unpurified cell-free expression lysate containing sc-antibodies

Epitope binning was tested on SPOC SPR biosensors for the first time using sensor-bound anti-HER2 VHH and anti-HER2 scFv (Trastuzumab) for interrogating HER2 epitopes. Recombinant HER2 ECD was first flowed across the sensor for binding to the HER2 VHH and HER2 scFv (first panel of Figure 10C-D). Next, commercial therapeutic antibodies with known epitopes were flowed across the sensor to determine binding patterns. As expected, Trastuzumab was able to bind to the HER2 VHH-HER2 ECD complex (Figure 10C) due to non-overlapping epitopes of Trastuzumab and HER2 VHH for HER2, but was not able to bind to the HER2 Trastuzumab scFv-HER2 ECD complex (Figure 10D), as the Trastuzumab scFv and Trastuzumab antibody recognize identical epitopes. Pertuzumab does not share epitopes with HER2 VHH nor Trastuzumab scFv (Figure 10A) and thus successfully bound both HER2 VHH-HER2 ECD and Trastuzumab scFv-HER2 ECD complexes, while CD20-binding Rituximab negative control bound neither (Figure 10C-D).

**Figure 10:**
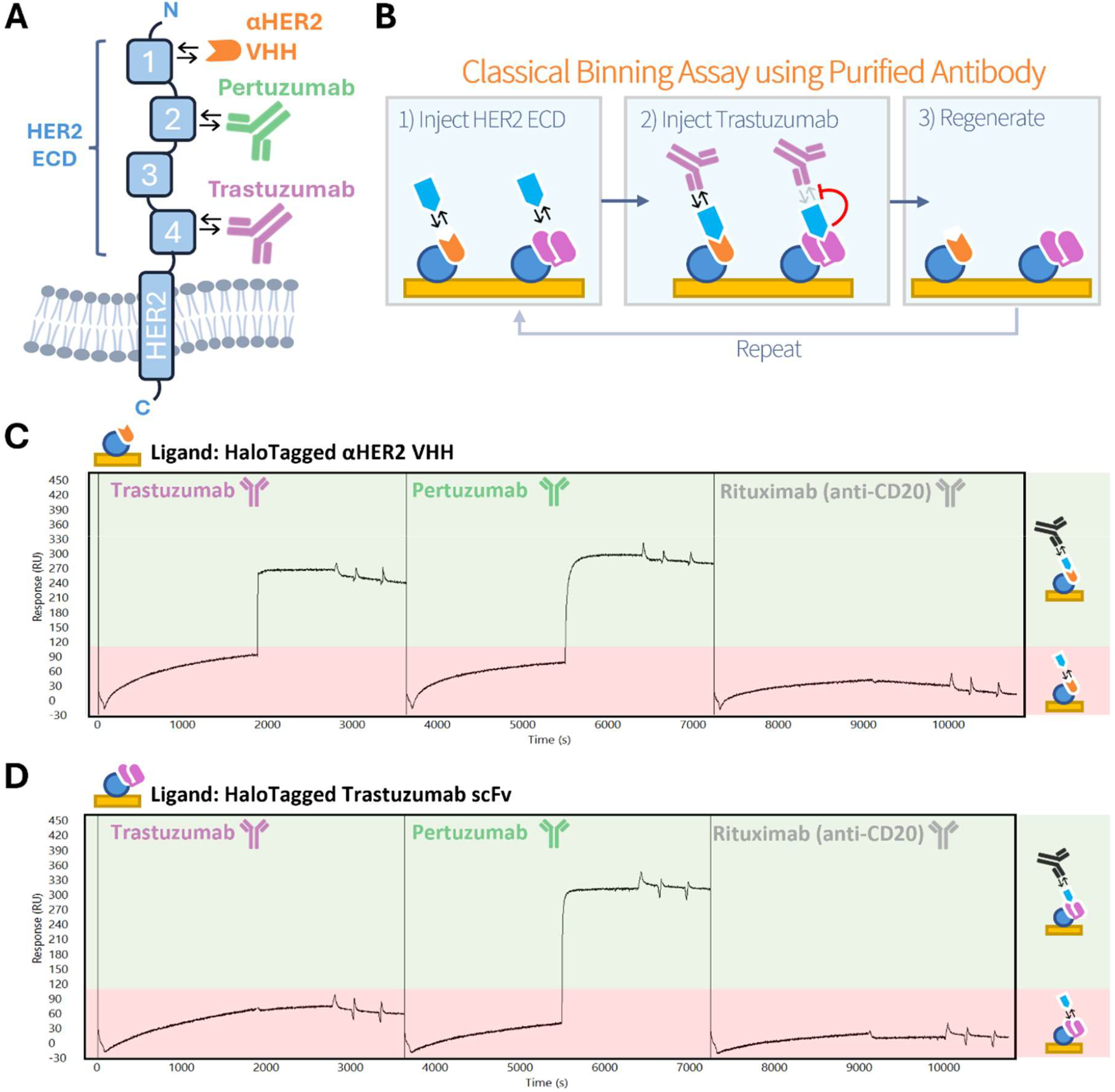
Demonstration of epitope binning using SPOC. (**A**) Schematic of HER2 depicting the four extracellular domains (ECD) and the non-overlapping domains targeted by the therapeutic antibodies and anti-HER2 VHH used in this study. The anti-HER2 VHH (orange) binds ECD 1, Pertuzumab (green) binds ECD 2, and Trastuzumab (pink) binds ECD 4. (**B**) Graphical depiction of the analyte injections involved in this binning experiment. Sensor bound HaloTagged (blue circle) anti-HER2 VHH (orange) and HaloTagged Trastuzumab scFv (pink) are depicted bound to the SPOC sensor surface. After HER2 ECD (light blue) injection and pre-binding, the sandwich antibody (Trastuzumab in this example) is injected and will bind any HER2 ECD bound to ligand with non-overlapping epitopes (i.e., anti-HER2 VHH). After antibody injection, the sensor is regenerated and HER2 ECD is re-injected and the cycle repeats for additional antibodies to be screened. (**C**) Sensorgram traces from sensor bound HaloTagged anti-HER2 VHH with injected Trastuzumab and Pertuzumab (green highlighted area) after HER2 ECD pre-loading (red space). (**D**) Sensorgram traces from sensor bound HaloTagged anti-HER2 Trastuzumab scFv binding HER2 ECD in the first injection, and then following Trastuzumab, Pertuzumab, or Rituximab injections.

Following validation of standard epitope binning procedures on SPOC SPR biosensors, we next tested if crude *in vitro* transcription/translation lysate (IVTT) from cell-free expression of scFv and VHH constructs could be used for epitope binning analytes in place of purified antibodies. After buffer exchange to remove non-protein small molecule components of the lysate (to avoid significant bulk shift on SPR), we repeated the epitope binning experiment after pre-binding with recombinant HER2 ECD (first panel of Figure 11B-C). As expected in these preliminary experiments, HER2 VHH lysate bound Trastuzumab scFv-HER2 ECD but not HER2 VHH-HER2 ECD complexes, and Trastuzumab scFv lysate bound HER2 VHH-HER2 ECD but not Trastuzumab scFv-HER2 ECD complexes (second panel of Figure 11B-C).

**Fig. 11:**
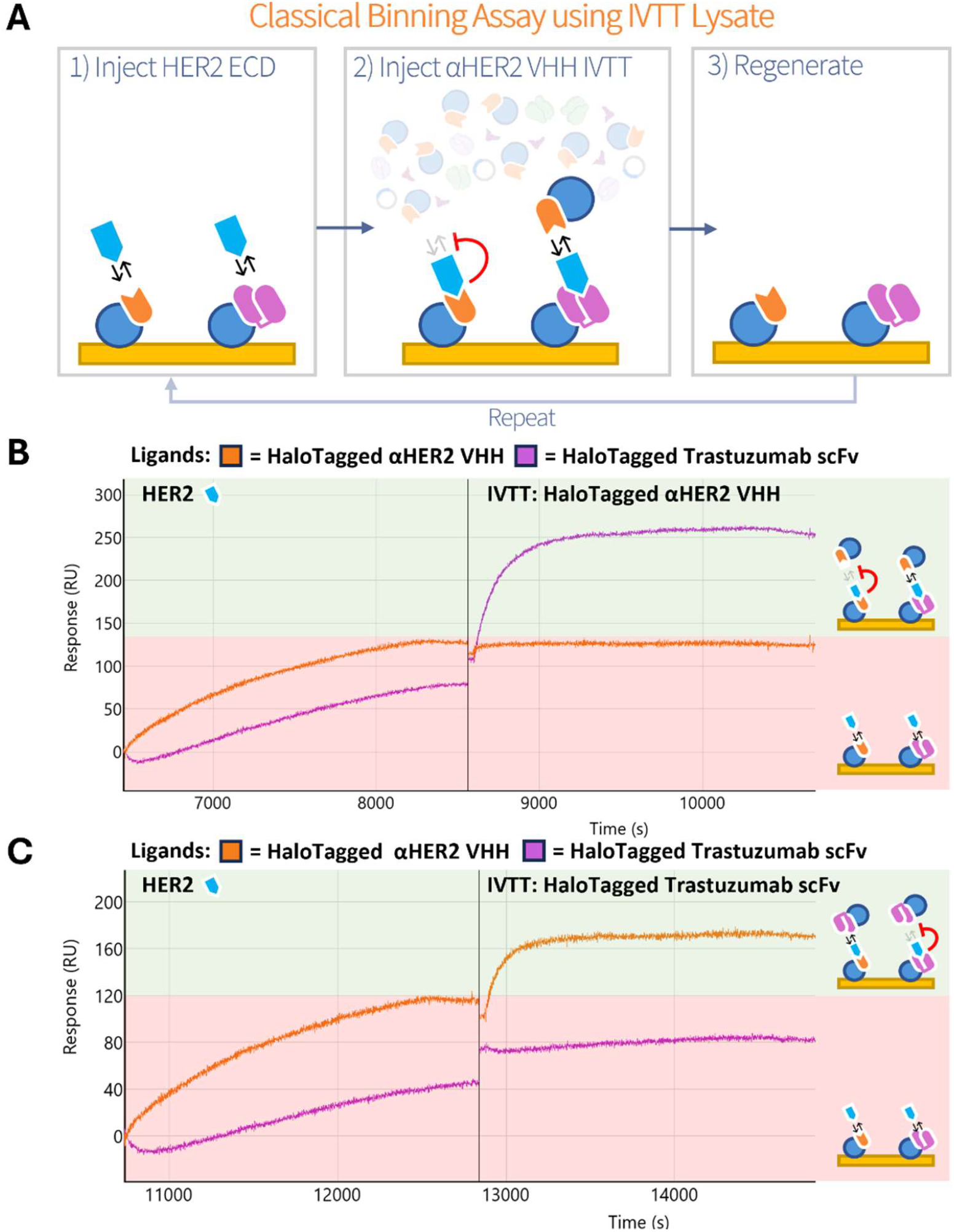
Demonstration of epitope binning using buffer-exchanged IVTT lysate as an antibody analyte source as opposed to purified antibody. (**A**) Graphical depiction of the analyte injections involved in this binning experiment. Sensor bound HaloTagged (blue circle) anti-HER2 VHH (orange) and HaloTagged Trastuzumab scFv (pink) are depicted bound to the SPOC sensor surface. After HER2 ECD (light blue) injection and pre-binding, diluted IVTT lysate containing free-floating HaloTagged anti-HER2 VHH or HaloTagged Trastuzumab scFv is injected to interrogate capability for productive sandwich formation, an indication of non-overlapping HER2 ECD binding sites with the corresponding ligand on the sensor surface. The sensor was regenerated and cycle repeated for subsequent HER2 ECD and IVTT lysate injections. (**B**) Injection of IVTT lysate expressing HaloTagged anti-HER2 VHH was performed after HER2 ECD preloading on a sensor containing both HaloTagged anti-HER2 VHH (orange trace) and HaloTagged Trastuzumab scFv (purple trace). Productive sandwich (green highlighted area) was only observed between IVTT anti-HER2 VHH and sensor bound HaloTagged Trastuzumab scFv. (**C**) Same experimental setup as in **B** but with IVTT HaloTagged Trastuzumab scFv sample used as analyte, only generating productive sandwich with sensor bound HaloTagged anti-HER2 VHH (green highlighted area) as expected.

### Design of single amino acid paratope mutations of HER2 VHH

Due to the clinical significance of HER2 therapeutic antibodies, the HER2 VHH was chosen for mutational analysis of the CDR regions that make up the paratope (antigen binding region) as a demonstration of SPOC biosensor utility in affinity maturation campaigns. Using data from Mitchell and Colwell^24^ for manual CDR assignment and automated confirmation via INDI^25^, CDR 1-3 were identified (Figure 12A) and each amino acid within the CDRs was individually mutated to alanine (substitution to a neutral, non-polar amino acid), aspartate (substitution to a negatively charged amino acid), lysine (substitution to a basic, positively charged amino acid), or serine (substitution to a neutral, polar amino acid), resulting in a total of 92 variants (Figure 12B). The DNA sequence was kept consistent except for mutations to ensure codon bias did not confound expression. The same codon was used within each amino acid substitution. Each variant was designed with a C-terminal HaloTag for covalent binding to the sensor.

**Figure 12:**
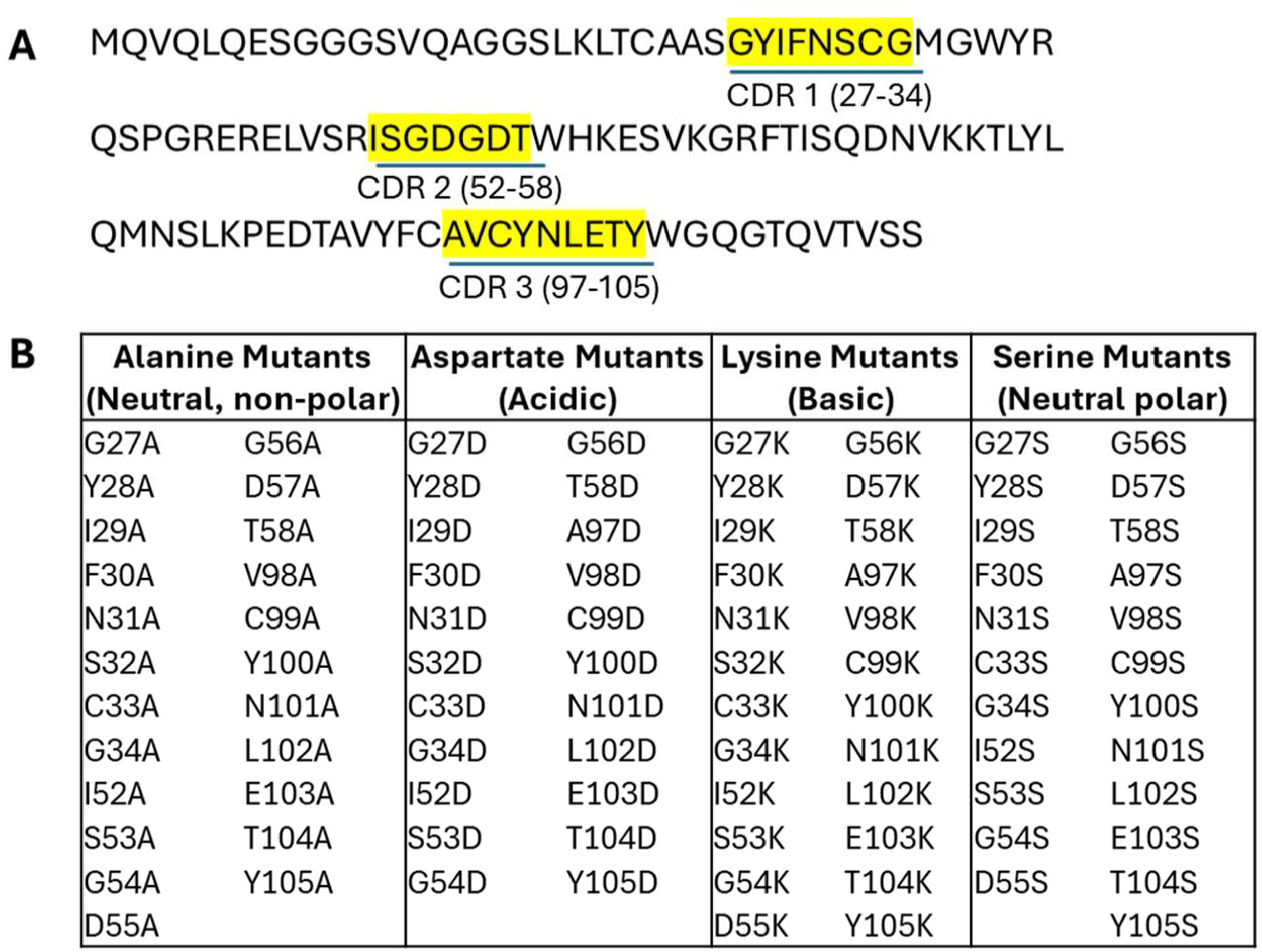
HER2 VHH (2Rs15d) wildtype sequence and resulting substitutions used in the mutation study. The sequence and identified CDRs of HER2 VHH (2Rs15d) (**A**). CDRs are highlighted in yellow and amino acid positions indicated for each. CDR amino acid substitutions to alanine, aspartate, lysine, and serine were designed into the constructs, resulting in 92 variants as shown in (**B**).

### Analysis of HER2 VHH CDR mutagenesis

SPOC biosensor chips with an sc-antibody library comprising mutated CDR variants were prepared from nanowell slides printed with DNA encoding constructs described above, and expressed using *E. coli* IVTT lysate.

Recombinant HER2 extracellular domain (ECD) was used as the analyte to measure binding properties of each mutant (Figure 13 and Supplementary Table 1). Calculation of affinity of wildtype/non-mutated HER2 VHH was measured to be 13.7 nM ± 0.745 nM from six individual spots. The averaged values from these spots were used for comparison to mutants (Figures 13-17). Substitution of S32K resulted in the lowest *K*_D_ at 31 nM ± 2.8 nM, with a >2-fold lower affinity than WT. Altering C33 or C99 to any amino acid ablated binding below the defined detection limit, suggesting these are critical paratope residues or provide structure required for binding via disulfide linkage; mutation of I52 to any of the four residues did the same. Interestingly, the reported crystal structure of this nanobody does not predict a disulfide linkage between these cysteine residues or others. Changing Y28 or T58 to any amino acid had no significant effect on the *K*_D_ compared to WT, while L102A, I29D, and L102S resulted in the highest affinity binders at 3.1 ± 0.85 nM, 3.1 nM ± 0.50 nM, and 3.6 nM ± 0.64 nM, respectively, all ∼4 fold higher than WT. These high affinity binders demonstrated non-canonical kinetic curves, where modest levels of binding were observed during the association phase, followed by a steep drop-off at the beginning of the dissociation phase and then a plateau where dissociation stalls. This may indicate a heterogeneous interaction between HER2 and the VHH for these mutants, where one form of interaction is highly stable. Overall, mutation to lysine appeared to be detrimental to binding affinity and mutation to aspartate obliterated or significantly reduced binding in more than a third of the mutated residues. Higher resolution sensorgrams are shown for WT and mutants C33A, N101D, S32K, G54S, and I29D (Figure 17).

**Figure 13:**
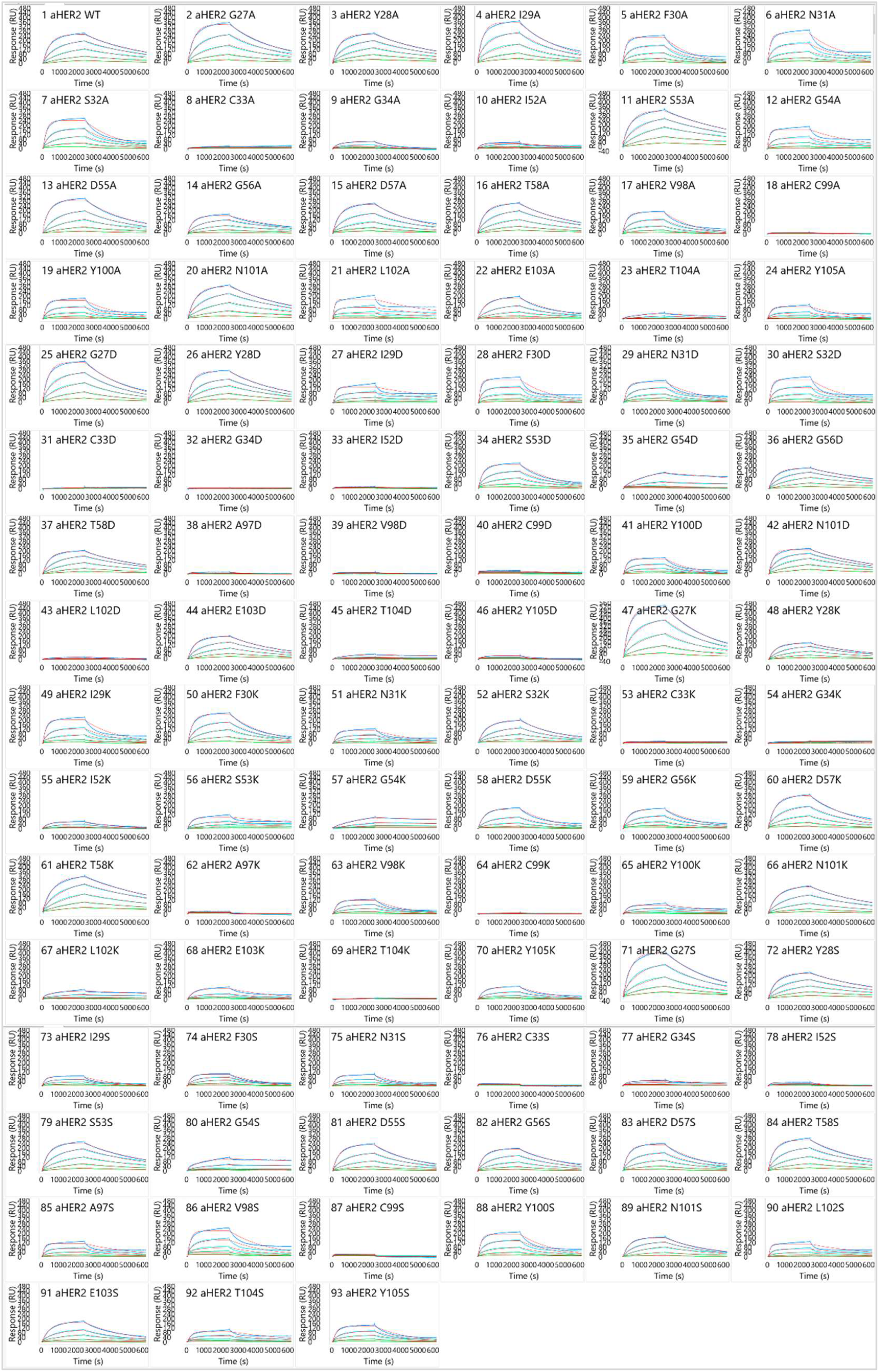
Sensorgrams showing HER2 ECD titration kinetics for a single replicate each of the 92 HER2 VHH mutants. Titration was performed using regenerative conditions with increasing concentrations of HER2 ECD (1.4 nM, 4.1 nM, 12 nM, 37 nM, 110 nM), with all injections overlayed onto a single sensorgram. The red lines indicate the kinetic model curve fits to underlying data plots.

**Figure 14:**
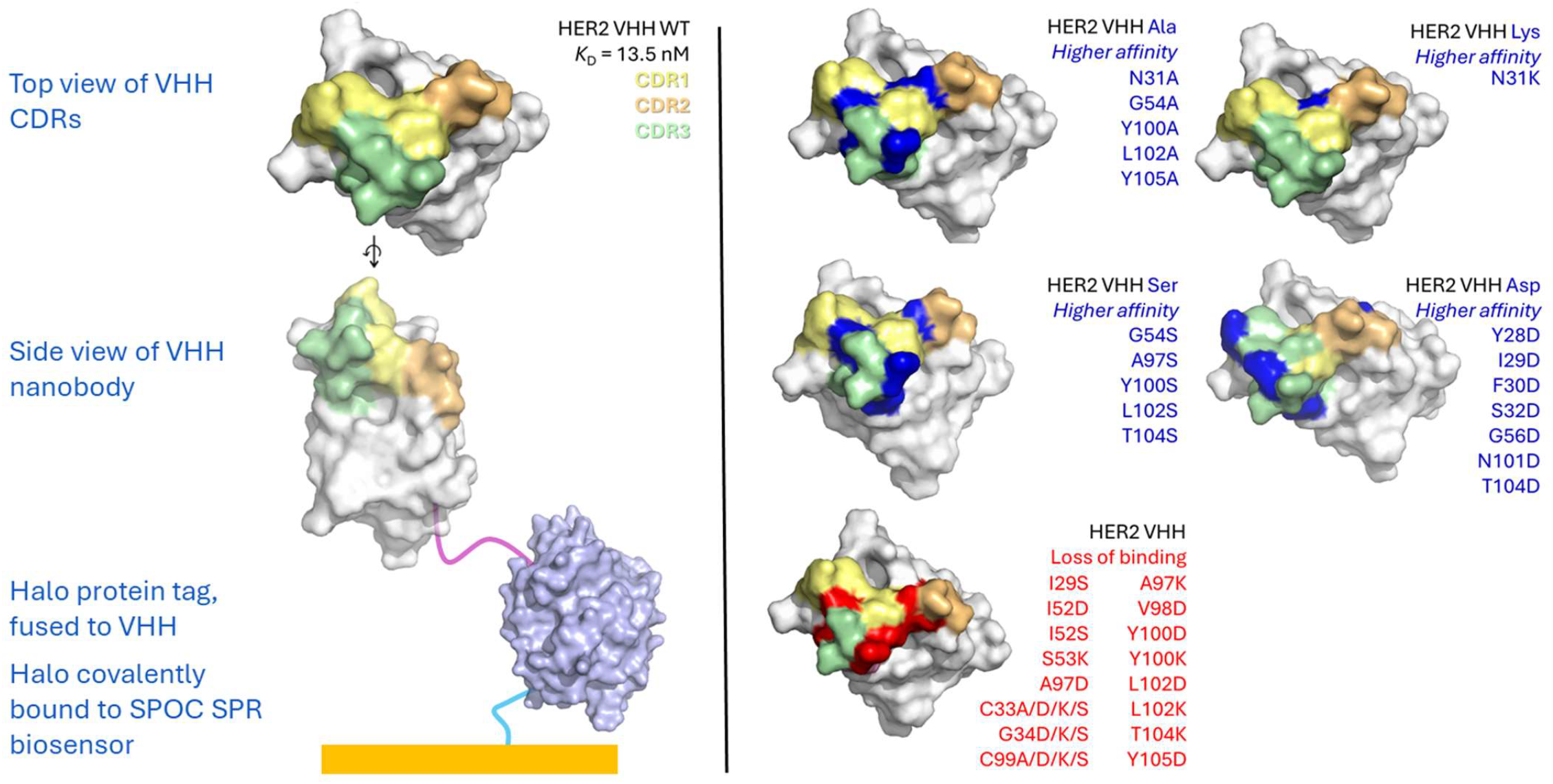
Visualization of residues that resulted in loss of binding or higher affinity upon substitution, in comparison to wildtype. CDRs 1, 2, and 3 are denoted in light yellow, light orange, and light green, respectively. Residues which produced higher affinity upon substitution are summarized visually in blue for each amino acid substitute (alanine, Ala; lysine, Lys; serine, Ser; aspartate, Asp) and specific substitutions are noted for each. Residues which resulted in loss of binding (red) or lower affinity 3-times or greater compared to wildtype (pink) are visualized in the lower left corner, with specific substitutions that resulted in loss of binding noted for each.

**Figure 15:**
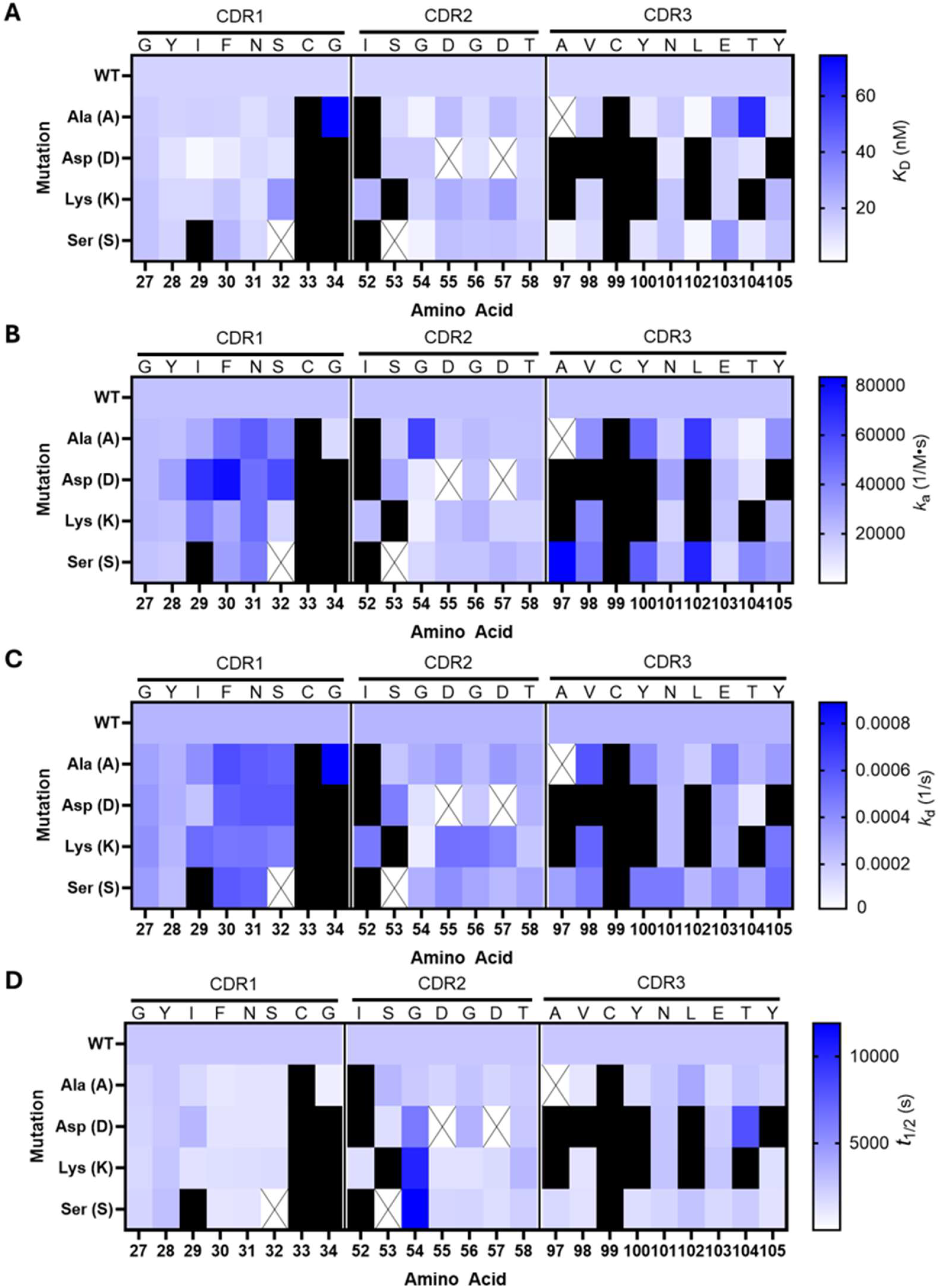
Heat maps visualizing kinetic binding data of the HER2 mutants. Affinity constant *K*D (**A**), on-rate *k*_a_ (**B**), off-rate *k*_d_ (**C**), and *t*_1/2_ (**D**) is shown. White boxes marked with X indicate the substitution is the same amino acid as wildtype and thus were not produced/duplicated. Black boxes indicate no binding was observed as previously defined in the methods. Bar graphs of this data are shown in **Figure 17**.

**Figure 16:**
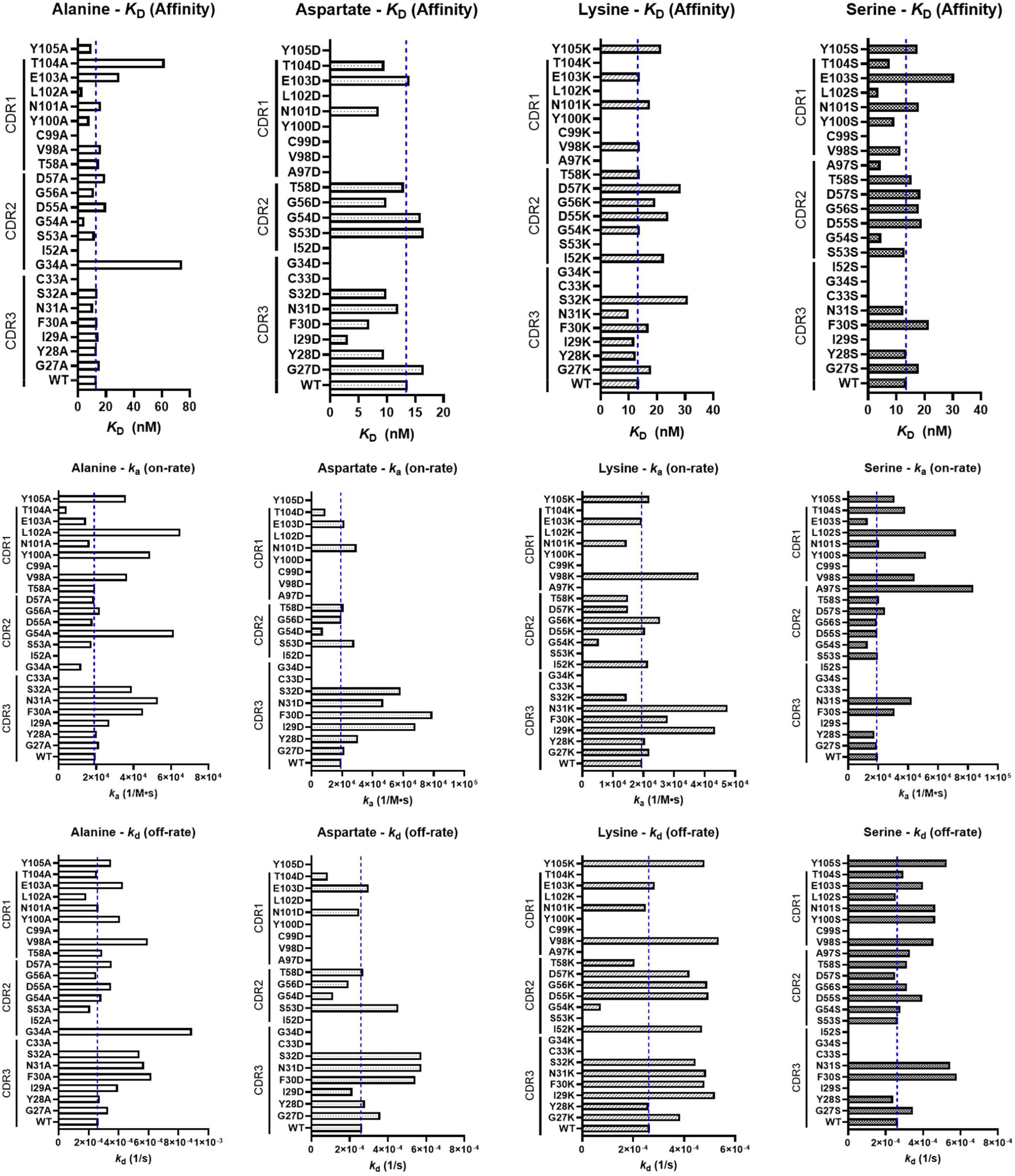
Bar graphs showing kinetic data obtained following a binding study of HER2 ECD against HER2 VHH mutants using a titration of HER2 ECD, organized by CDR and kinetic parameter. Affinity constant *K*_D_ for each amino acid substitution is shown in the top row, on-rate *k*_a_ is shown in the middle row, and off-rate *k*_d_ is shown in the bottom row. Missing bars indicate no binding was observed as defined in the methods.

**Figure 17:**
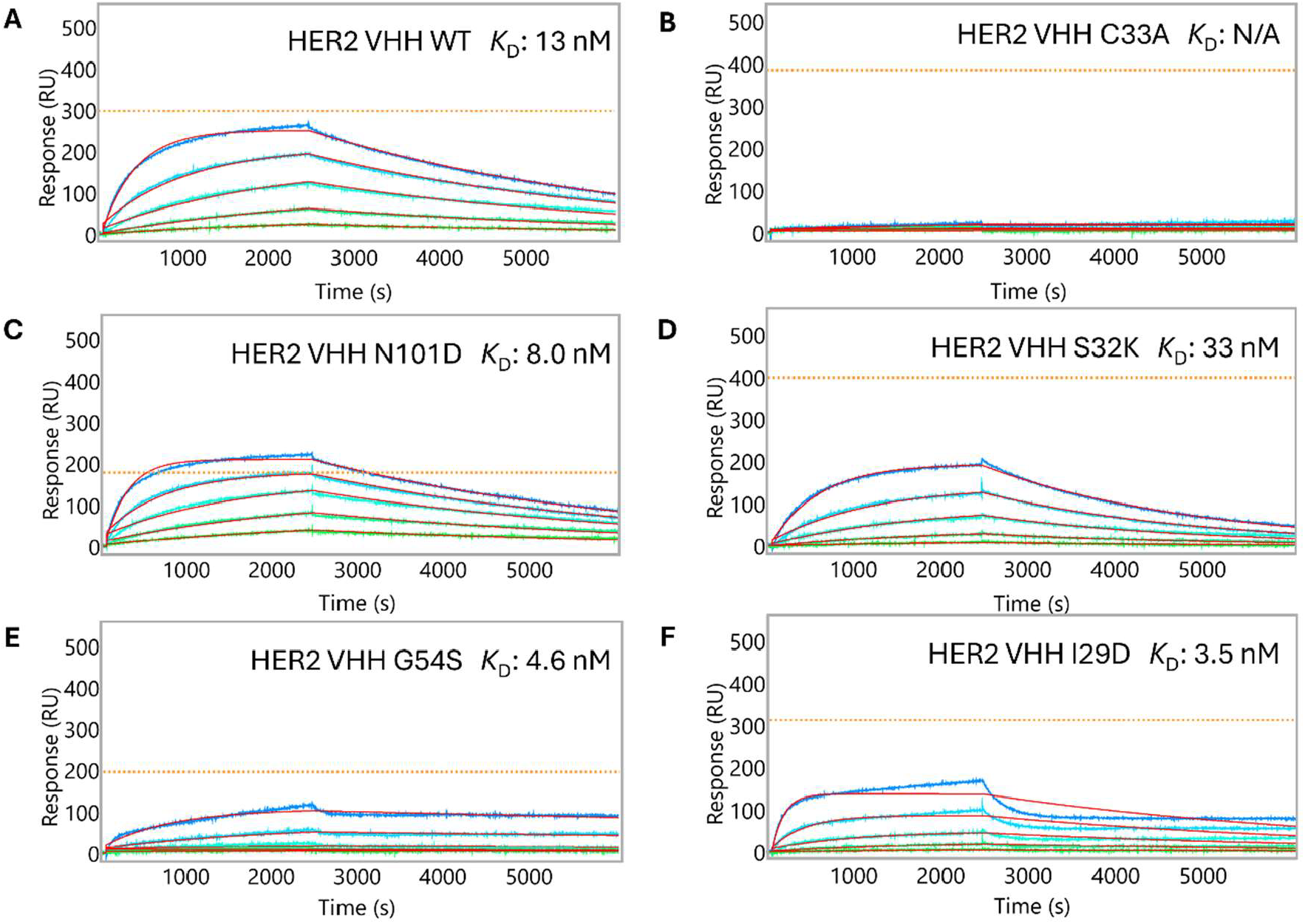
Select sensorgrams showing mutants with higher affinity compared to WT sequence. (A) WT, (B) loss of binding with C33A mutation, (C) higher affinity binding paired with higher overall binding with N101D mutation, (D) lowest affinity binder with D55K mutation, (E) high affinity with little loss binding, though with lower overall binding with G54S, and (F) high affinity binding with non-canonical kinetic curvature. Affinity curves (red) fitted to entire curve; Right: Curves fitted only for the dissociation (red). Red line: fitted kinetic curves. Orange dotted line: level of VHH bound to sensor, defined by the maximum signal (RU) of anti-HaloTag antibody binding used for capture validation.

### Analysis of affinity of HER2 ECD for HER2 VHH produced from a titration of DNA encoding HER2 VHH

To determine whether evaluated kinetic parameters were dependent on the quantitative amounts of the mutated protein spots across the SPR chip, as each of the protein mutants may express at different levels, we performed an experiment where we titrated the printed amount of DNA encoding WT HER2 VHH into nanowells. We observed that although the magnitude of WT HER2 ECD binding signal increased with printed DNA concentration, the evaluated affinity remained effectively the same. This shows that affinity is consistent and independent of levels or variation in protein quantities captured at each spot on the SPR biosensors. Affinity values for HER2 ECD binding to HER2 VHH (wildtype) expressed from DNA printed at 11, 25, 33, 50, 75, and 100 ng/μL were measured to be 14 nM, 15 nM, 15 nM, 17 nM, 15 nM, and 18 nM, respectively, for an average of 16 nM ± 1.4 nM (Figure 18).

**Figure 18:**
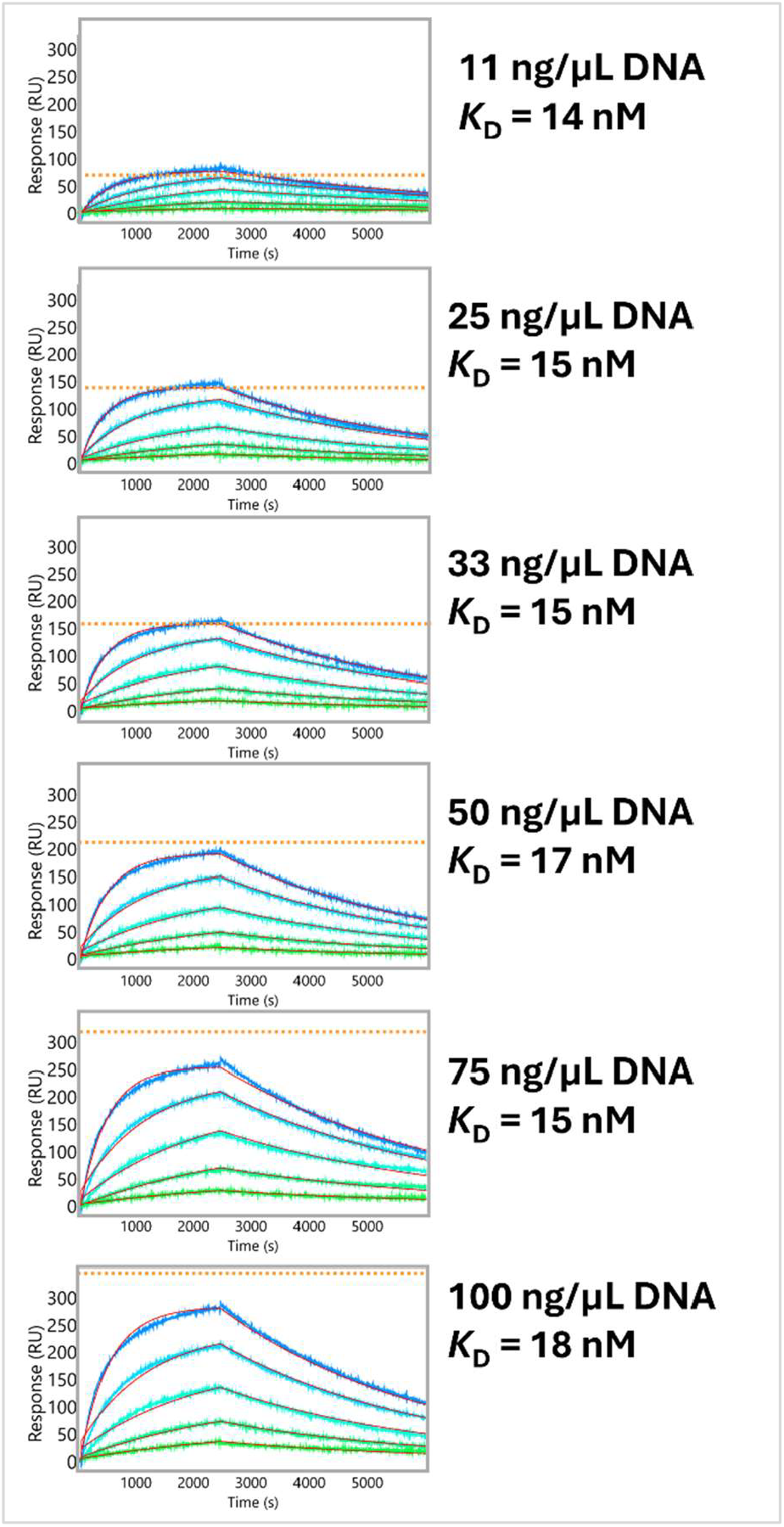
Varying input DNA concentration results in similar affinities for expressed protein. DNA encoding HER2 VHH wildtype was titrated during the nanowell printing process and the resulting protein expressed and captured onto the SPOC sensor was evaluated for binding with HER2 ECD. Red line: fitted kinetic curves. Orange dotted line: level of VHH bound to sensor, defined by the maximum signal (RU) of anti-HaloTag antibody binding used for capture validation.

## Discussion

We have previously described a highly multiplexed method for measuring the kinetics of hundreds to thousands of protein interactions on a single biosensor using cell-free expression and high throughput SPR^12^. Here, we describe a new application of the SPOC platform demonstrating the production and testing of single chain antibodies and Fabs via *in situ* cell-free expression and capture-purification onto SPR biosensors, with a use-case application of mutational scanning of an anti-HER2 VHH paratope.

For the development of this application, fluorescence-based assays were used as an initial test to validate antibody-target binding due to the ability to control a matrix of factors at one time for optimization of dilutions, buffers, and choice of detection antibody with high sensitivity. Following this, a set of sc-antibody molecules were tested using a SPOC chip on a custom Carterra SPR instrument to determine if scFv and VHH constructs produced in cell-free systems are functional and bind to respective targets with high specificity. This facilitates characterizing on-chip sc-antibody selectivity with significantly higher throughput, and to collect kinetic measurements of antigen binding for in-depth characterization and affinity ranking.

This paper demonstrates that the SPOC Protein Nano Factory (PNF) system produces functional Fabs and single chain antibodies in the two forms tested directly on SPR biosensors: scFvs derived from full-length antibodies and single-domain antibody VHH, also commonly known as nanobodies. These proteins bound highly detectable target antigen on SPOC chips via SPR (Figure 7-8). All constructs were synthesized and their expression products were tested on SPOC biosensors. Expressed protein was confirmed to be present on the sensor at readily detectable levels as measured via anti-HaloTag antibody. It was immediately clear that 5 scFvs (targeting IFNγ, HSA, EGFR, HER2, and CEA) and 2 VHH (targeting TNFa and HER2) bound their label-free target antigen at high levels at first assay with no additional optimization of SPR or expression parameters required. These constructs correlated well with previously reported affinity, with these immediately detectable antigens binding to constructs with low to sub-nanomolar affinity.

The constructs tested here had sequences taken directly from literature and were converted to scFvs using a single common linker of (Gly_4_Ser)_3_, without experimenting with multiple linker formats or other scFv components or scaffolds. We have also tested CA15.3, TNFa, and other HER2 scFv constructs using this common linker. CA15.3 and TNFa scFv have shown no binding thus far in this single-shot testing, while IL-6 and p53 easily bound their target, though specificity data is pending. Traditionally, scFvs are designed in the format of V_H_-(Gly_4_Ser)_3_-V_L_, as the most characterized and preferred linker format. However, given the precise scaffolding and folding requirements of scFvs, quite often this requires further optimization with different linker scaffolds if the initial format results in misfolding or aggregation, with no binding of target. Researchers often evaluate different linker lengths, different linker types, switching the order of the V_H_ and V_L_, moving a purification or expression tag from one terminus to another, etc. We plan to test any non-functional scFv constructs with alternative linker designs and other formats in the future.

Given the success of scFv constructs, we created matching Fab constructs for testing on SPOC SPR biosensors. Constant regions for the heavy and light chain were the same as reported for full length antibodies in literature with the exception of the CEA Fab. The CEA antibody fragments used in this study were originally derived from a scFv library. To overcome this, constant regions from Emapalumab (anti-IFNγ) were used for the CEA Fab. The HaloTag was expressed as part of the heavy chain fragment on the C-terminus, and the light chain was expressed without a HaloTag. Gene fragments encoding the heavy chain and the light chain were co-printed into the same nanowell and were thus expressed and assembled within the same nanowell, resulting in an assembled Fab on the SPOC biosensor. Binding of the Fab targets to target analytes were successfully detected and found to be selective (Figure 8).

From the scFv and VHH constructs, two VHH and one scFv were further analyzed to calculate affinity and limit of detection via injection of a titration of antigen. For the TNFα VHH, an approximate affinity of 149 pM was calculated, nearly 4-fold higher than the reported affinity of 540 pM for this VHH^17^. The affinity measurements for this study were collected using a 1 hour association and 4 hour dissociation time to accurately analyze kinetics of binders with affinity in the picomolar range. Additionally, when using SPR biosensing to characterize kinetic parameters of very high affinity binders (picomolar to femtomolar affinities), we also have the option of using the “chaser assay” method to accurately characterize ultra-low-dissociation-rate binding events measured by SPR instrument on SPOC chips. Used in the field of enzymology and more recently described in detail for application to SPR biosensing methodology by Quinn et al^26^, this method uses a “chaser probe” to measure the fraction of free binding sites on the ligand epitope of interest after an acceptable dissociation time (minutes-hours versus many hours-days), resulting in high resolution readout at very high affinity. For the HER2 VHH, the measured affinity for recombinant HER2 ECD was calculated to be 13.5 nM, very close to the previously reported affinity for this VHH of 4 nM^18^, while the Trastuzumab scFv affinity for HER2 ECD was calculated to be 6.7 nM. The reported affinity of Trastuzumab according to FDA documentation is 5 nM.

As demonstration for use of the SPOC technology in drug development, we created a panel of paratope mutants of the HER2 VHH and measured the binding characteristics of HER2 to each mutant simultaneously on a single chip for direct comparison to one another. By mutating each amino acid individually to alanine within the three complementarity determining regions, it can be determined which amino acid side chains are critical for antigen binding. Similarly, by mutating the same residues individually to aspartate (acidic), lysine (basic), or serine (polar), we can determine if changing the side chain properties improves or reduces antigen binding. We found that several mutants improved overall affinity, though overall binding levels appeared to suffer as a result. This may be considered an acceptable tradeoff, however. Nearly all improvements in affinity were made by substitution to either Asp or Ser. Mutation of G54 resulted in equal or higher affinity binding, indicating this particular residue may be mutated for further improvement of *k*^D^. Using data-trained AI-driven models for iterative design on more optimal sequences, 10 – 100x improvements in affinity via modulation of critical amino acids becomes possible.

When using AI models for sc-antibody engineering, the SPOC platform addresses critical bottlenecks in the “build” and “test” phases by enabling the interrogation of thousands of nanobody candidates in a single assay. This facilitates the down-selection of the most promising candidates based on objective, high-affinity-resolution kinetic data. SPOC supports affinity maturation cycles by facilitating detailed characterization and engineering of antibody paratopes, enabling the design of variants with improved affinity, specificity, and selectivity. With its relatively low production cost, SPOC allows for comprehensive mutational scanning of antibody CDRs, substituting each position with all other amino acids (or a select subset) on a single SPR chip. This approach generates a complete, amino acid-level kinetic dataset to inform the “learn” phase and subsequent iterations of design-build-test-learn (DBTL) cycles. By performing this analysis on the same chip, SPOC ensures direct and confident comparisons of measurements between mutants and the wild-type baseline. Kinetic parameter ranking then enables the selection of binders tailored to specific therapeutic modalities or mechanisms of interest, such as those that bind with highest affinities, or that bind quickly (fast on-rate) at high levels, or that bind very tightly (slow off-rate), or a combination of these.

In conclusion, we have described for the first time a method for producing single chain antibodies and Fabs on SPOC SPR biosensor chips via cell-free protein expression in a highly multiplexed format for label-free detection and characterization of target antigen binding. This method allows the user to simply use a DNA input for biosensor production of thousands of unique single chain antibody constructs, rather than expressing each construct of interest individually at in sufficient amounts followed by spotting or capturing onto a SPR biosensor chip for kinetic assay. SPOC can be applied to evaluating the affinities of antibodies produced via computational modeling to reduce downstream costs and increase the throughput of testing. For example, we demonstrated expression/production of a panel of 92 mutationally scanned variants from the CDR of HER2 VHH on one SPR biosensor, directly from DNA in an *E. coli* IVTT lysate. The SPOC SPR biosensor, meanwhile, has the capacity to interrogate up to 2,400 mutants, as reported previously. The result is the capability to collect and measure kinetic data at-scale at significantly lower cost per sequence than traditional recombinant production methods. This demonstrates a significant improvement to the ‘build’ and ‘test’ cycles required in traditional drug discovery, AI-enabled, and AI-driven workflows. Lastly, we demonstrated how a lead candidate can be mutationally scanned at low-cost and at high-throughput with kinetic measurements to support deep paratope characterization for subsequent affinity maturation campaigns, proposing the application of SPOC platform for iterative DBTL cycles to support traditional and AI drug discovery pipelines. The future goal for this application will be to fully incorporate SPOC analysis into AI drug discovery workstreams, to demonstrate how the improved ‘build’ and ‘test’ phases with increased sequence diversity testing capabilities, will improve the subsequent iterative ‘learn’ and ‘design’ phases and result in improved lead drug candidates, ultimately towards improving drug success rates.

## Acknowledgements

We acknowledge partial funding support from NIH SBIR grant 1R44TR004297. We acknowledge Sharrol Bachas for helpful discussions on selecting the four diverse amino acids for CDR mutational scanning substitutions.

