## Supplementary Data for "An approach to produce thousands of single-chain antibody variants on a SPR biosensor chip for measuring target binding kinetics and for deep characterization of antibody paratopes"

Supplementary Table 1: Analysis of HER2 VHH mutation kinetics using 1:1 binding model, **Replicate 1**

| Name | $k_a$ (1/M·s) | $k_d$ (1/s) | *KD* (nM) | $t_{1/2}$ (s) | Rmax (expected) | Rmax @ 110 nM HER2 | HaloTag Rmax @ 130 nM |
|---|---|---|---|---|---|---|---|
| aHER2 WT | 2.0E+04 | 2.6E-04 | 13 nM | 2.6E+03 | 507.4 | 261.3 | 303.6 |
| aHER2 G27A | 2.2E+04 | 3.3E-04 | 15 nM | 2.1E+03 | 521.4 | 351.7 | 431.6 |
| aHER2 Y28A | 1.9E+04 | 2.5E-04 | 13 nM | 2.8E+03 | 633.3 | 258.3 | 245.3 |
| aHER2 I29A | 2.6E+04 | 3.9E-04 | 15 nM | 1.8E+03 | 655.2 | 362.9 | 398.2 |
| aHER2 F30A | 4.6E+04 | 6.3E-04 | 14 nM | 1.1E+03 | 183.2 | 241.6 | 300.9 |
| aHER2 N31A | 5.2E+04 | 4.9E-04 | 9.5 nM | 1.4E+03 | 137.4 | 284.9 | 530.1 |
| aHER2 S32A | 4.3E+04 | 5.4E-04 | 13 nM | 1.3E+03 | 118.1 | 260.5 | 461.4 |
| aHER2 C33A | N/A | N/A | N/A | N/A | N/A | 22.9 | 491.9 |
| aHER2 G34A | 1.1E+04 | 1.0E-03 | 90 nM | 6.9E+02 | 238.8 | 62.7 | 371.5 |
| aHER2 I52A | N/A | N/A | N/A | N/A | N/A | 57.2 | 633.4 |
| aHER2 S53A | 1.7E+04 | 2.0E-04 | 12 nM | 3.5E+03 | 764.6 | 326.0 | 471.4 |
| aHER2 G54A | 8.0E+04 | 3.3E-04 | 4.1 nM | 2.1E+03 | 27.0 | 190.9 | 328.3 |
| aHER2 D55A | 2.0E+04 | 3.5E-04 | 18 nM | 2.0E+03 | 372.4 | 302.7 | 469.4 |
| aHER2 G56A | 2.6E+04 | 2.6E-04 | 10 nM | 2.6E+03 | 147.5 | 170.3 | 328.2 |
| aHER2 D57A | 1.8E+04 | 3.5E-04 | 20 nM | 2.0E+03 | 462.3 | 261.6 | 320.9 |
| aHER2 T58A | 1.8E+04 | 2.9E-04 | 16 nM | 2.4E+03 | 588.6 | 262.9 | 305.8 |
| aHER2 V98A | 3.5E+04 | 6.0E-04 | 17 nM | 1.2E+03 | 203.2 | 194.9 | 231.7 |
| aHER2 C99A | N/A | N/A | N/A | N/A | N/A | 12.1 | 489.2 |
| aHER2 Y100A | 5.3E+04 | 4.8E-04 | 9.1 nM | 1.4E+03 | 82.9 | 179.1 | 285.9 |
| aHER2 N101A | 1.7E+04 | 2.5E-04 | 15 nM | 2.8E+03 | 511.0 | 291.1 | 478.6 |
| aHER2 L102A | 9.4E+04 | 2.3E-04 | 2.5 nM | 3.0E+03 | 24.2 | 203.0 | 289.6 |
| aHER2 E103A | 1.6E+04 | 4.3E-04 | 27 nM | 1.6E+03 | 129.9 | 188.9 | 405.1 |
| aHER2 T104A | 4.4E+03 | 2.6E-04 | 60 nM | 2.6E+03 | 92.8 | 51.3 | 467.0 |
| aHER2 Y105A | 4.2E+04 | 4.5E-04 | 11 nM | 1.5E+03 | 12.8 | 121.1 | 402.6 |
| aHER2 G27D | 2.3E+04 | 3.7E-04 | 16 nM | 1.9E+03 | 688.6 | 356.8 | 388.2 |
| aHER2 Y28D | 2.8E+04 | 2.9E-04 | 11 nM | 2.4E+03 | 485.1 | 280.4 | 240.1 |
| aHER2 I29D | 6.5E+04 | 2.3E-04 | 3.5 nM | 3.1E+03 | 29.6 | 166.3 | 310.1 |
| aHER2 F30D | 7.7E+04 | 5.3E-04 | 6.9 nM | 1.3E+03 | 105.8 | 223.3 | 274.1 |
| aHER2 N31D | 4.2E+04 | 4.8E-04 | 11 nM | 1.5E+03 | 200.8 | 191.2 | 224.3 |
| aHER2 S32D | 6.3E+04 | 4.9E-04 | 7.8 nM | 1.4E+03 | 78.5 | 227.9 | 322.2 |
| aHER2 C33D | N/A | N/A | N/A | N/A | N/A | 7.1 | 353.6 |
| aHER2 G34D | N/A | N/A | N/A | N/A | N/A | 3.8 | 303.1 |
| aHER2 I52D | N/A | N/A | N/A | N/A | N/A | 15.8 | 437.0 |
| aHER2 S53D | 3.0E+04 | 4.8E-04 | 16 nM | 1.4E+03 | 138.5 | 217.1 | 235.3 |
| aHER2 G54D | 6.3E+03 | 9.8E-05 | 16 nM | 7.1E+03 | 82.0 | 139.0 | 315.4 |
| aHER2 G56D | 1.9E+04 | 2.1E-04 | 11 nM | 3.2E+03 | 327.9 | 178.8 | 280.5 |
| aHER2 T58D | 2.1E+04 | 2.6E-04 | 13 nM | 2.6E+03 | 412.0 | 200.2 | 255.5 |
| aHER2 A97D | N/A | N/A | N/A | N/A | N/A | 15.0 | 475.1 |
| aHER2 V98D | N/A | N/A | N/A | N/A | N/A | 12.1 | 501.9 |
| aHER2 C99D | N/A | N/A | N/A | N/A | N/A | 28.8 | 311.2 |
| aHER2 Y100D | N/A | N/A | N/A | N/A | N/A | 138.9 | 142.2 |
| aHER2 N101D | 3.2E+04 | 2.5E-04 | 8.0 nM | 2.7E+03 | 496.1 | 220.3 | 185.9 |
| aHER2 L102D | N/A | N/A | N/A | N/A | N/A | 19.4 | 149.4 |
| aHER2 E103D | 1.9E+04 | 3.1E-04 | 16 nM | 2.3E+03 | 406.8 | 195.7 | 206.9 |
| aHER2 T104D | 9.3E+03 | 1.0E-04 | 11 nM | 6.8E+03 | 15.4 | 44.7 | 195.5 |

| | | | | | | | |
|---|---|---|---|---|---|---|---|
| aHER2 Y105D | N/A | N/A | N/A | N/A | N/A | 32.3 | 423.1 |
| aHER2 G27K | 2.3E+04 | 3.6E-04 | 16 nM | 1.9E+03 | 801.9 | 531.7 | 751.2 |
| aHER2 Y28K | 1.9E+04 | 2.7E-04 | 14 nM | 2.6E+03 | 301.7 | 143.0 | 146.5 |
| aHER2 I29K | 4.7E+04 | 5.3E-04 | 11 nM | 1.3E+03 | 75.5 | 222.6 | 506.9 |
| aHER2 F30K | 2.8E+04 | 4.8E-04 | 17 nM | 1.4E+03 | 257.3 | 259.8 | 294.4 |
| aHER2 N31K | 4.5E+04 | 4.2E-04 | 9.2 nM | 1.7E+03 | 22.7 | 123.0 | 237.4 |
| aHER2 S32K | 1.2E+04 | 4.1E-04 | 33 nM | 1.7E+03 | 265.5 | 190.2 | 398.2 |
| aHER2 C33K | N/A | N/A | N/A | N/A | N/A | 17.0 | 288.9 |
| aHER2 G34K | N/A | N/A | N/A | N/A | N/A | 15.5 | 419.2 |
| aHER2 I52K | 2.0E+04 | 5.3E-04 | 27 nM | 1.3E+03 | 124.2 | 61.0 | 389.6 |
| aHER2 S53K | N/A | N/A | N/A | N/A | N/A | 116.9 | 357.1 |
| aHER2 G54K | 3.5E+03 | 5.4E-05 | 15 nM | 1.3E+04 | 134.1 | 87.9 | 262.7 |
| aHER2 D55K | 2.0E+04 | 5.3E-04 | 26 nM | 1.3E+03 | 146.2 | 178.5 | 405.8 |
| aHER2 G56K | 2.5E+04 | 4.9E-04 | 20 nM | 1.4E+03 | 190.3 | 174.5 | 557.9 |
| aHER2 D57K | 1.8E+04 | 4.0E-04 | 22 nM | 1.7E+03 | 349.0 | 291.1 | 405.1 |
| aHER2 T58K | 1.4E+04 | 2.0E-04 | 15 nM | 3.4E+03 | 1120.8 | 314.7 | 423.9 |
| aHER2 A97K | N/A | N/A | N/A | N/A | N/A | 21.9 | 375.4 |
| aHER2 V98K | 3.8E+04 | 5.6E-04 | 15 nM | 1.2E+03 | 53.2 | 126.5 | 325.6 |
| aHER2 C99K | N/A | N/A | N/A | N/A | N/A | 5.1 | 398.9 |
| aHER2 Y100K | N/A | N/A | N/A | N/A | N/A | 89.3 | 239.2 |
| aHER2 N101K | 1.4E+04 | 2.6E-04 | 19 nM | 2.7E+03 | 352.2 | 238.9 | 366.4 |
| aHER2 L102K | N/A | N/A | N/A | N/A | N/A | 82.7 | 208.0 |
| aHER2 E103K | 2.3E+04 | 3.7E-04 | 16 nM | 1.9E+03 | 55.8 | 103.3 | 342.1 |
| aHER2 T104K | N/A | N/A | N/A | N/A | N/A | 7.5 | 317.2 |
| aHER2 Y105K | 2.4E+04 | 5.2E-04 | 22 nM | 1.3E+03 | 85.4 | 109.6 | 303.5 |
| aHER2 G27S | 1.8E+04 | 3.3E-04 | 18 nM | 2.1E+03 | 1029.8 | 403.7 | 514.8 |
| aHER2 Y28S | 1.7E+04 | 2.2E-04 | 13 nM | 3.1E+03 | 561.0 | 225.5 | 248.8 |
| aHER2 I29S | N/A | N/A | N/A | N/A | N/A | 85.2 | 98.3 |
| aHER2 F30S | 4.1E+04 | 5.4E-04 | 13 nM | 1.3E+03 | 75.9 | 105.9 | 86.5 |
| aHER2 N31S | 4.6E+04 | 6.1E-04 | 13 nM | 1.1E+03 | 19.5 | 97.1 | 139.3 |
| aHER2 C33S | N/A | N/A | N/A | N/A | N/A | 14.4 | 300.0 |
| aHER2 G34S | N/A | N/A | N/A | N/A | N/A | 51.8 | 286.7 |
| aHER2 I52S | N/A | N/A | N/A | N/A | N/A | 29.0 | 430.6 |
| aHER2 S53S | 2.0E+04 | 2.6E-04 | 13 nM | 2.7E+03 | 470.9 | 252.9 | 265.0 |
| aHER2 G54S | 1.1E+04 | 4.9E-05 | 4.6 nM | 1.4E+04 | 80.7 | 114.7 | 203.0 |
| aHER2 D55S | 1.9E+04 | 3.9E-04 | 20 nM | 1.8E+03 | 276.3 | 234.3 | 312.5 |
| aHER2 G56S | 2.1E+04 | 3.6E-04 | 17 nM | 1.9E+03 | 255.5 | 196.6 | 347.2 |
| aHER2 D57S | 2.4E+04 | 4.3E-04 | 18 nM | 1.6E+03 | 192.4 | 238.6 | 354.5 |
| aHER2 T58S | 2.2E+04 | 2.9E-04 | 13 nM | 2.4E+03 | 449.5 | 281.2 | 351.1 |
| aHER2 A97S | 7.3E+04 | 3.3E-04 | 4.4 nM | 2.1E+03 | 13.5 | 129.9 | 189.1 |
| aHER2 V98S | 4.8E+04 | 4.3E-04 | 9.0 nM | 1.6E+03 | 138.0 | 246.4 | 610.3 |
| aHER2 C99S | N/A | N/A | N/A | N/A | N/A | 15.1 | 520.2 |
| aHER2 Y100S | 4.8E+04 | 5.1E-04 | 11 nM | 1.4E+03 | 146.3 | 210.1 | 301.2 |
| aHER2 N101S | 2.1E+04 | 3.7E-04 | 18 nM | 1.9E+03 | 148.6 | 166.5 | 276.9 |
| aHER2 L102S | 8.9E+04 | 2.8E-04 | 3.2 nM | 2.5E+03 | 13.1 | 126.9 | 173.0 |
| aHER2 E103S | 1.5E+04 | 4.2E-04 | 27 nM | 1.7E+03 | 50.1 | 176.7 | 328.6 |
| aHER2 T104S | 4.0E+04 | 2.9E-04 | 7.2 nM | 2.4E+03 | 34.2 | 102.4 | 295.4 |
| aHER2 Y105S | 3.6E+04 | 5.4E-04 | 15 nM | 1.3E+03 | 45.0 | 143.4 | 370.0 |

Supplementary Table 1: Analysis of HER2 VHH mutation kinetics using 1:1 binding model, **Replicate 2**

| Name | $k_a$ (1/M·s) | $k_d$ (1/s) | *KD* (nM) | $t_{1/2}$ (s) | Rmax (expected) | Rmax @ 110 nM HER2 | HaloTag Rmax @ 130 nM |
|---|---|---|---|---|---|---|---|
| aHER2 WT | 1.9E+04 | 2.7E-04 | 14 nM | 2.5E+03 | 415.9 | 241.7 | 285.1 |
| aHER2 G27A | 2.1E+04 | 3.3E-04 | 16 nM | 2.1E+03 | 580.5 | 395.6 | 503.1 |
| aHER2 Y28A | 2.2E+04 | 3.0E-04 | 14 nM | 2.3E+03 | 519.2 | 263.0 | 262.6 |
| aHER2 I29A | 2.8E+04 | 4.0E-04 | 14 nM | 1.7E+03 | 411.4 | 298.1 | 328.1 |
| aHER2 F30A | 4.4E+04 | 6.1E-04 | 14 nM | 1.1E+03 | 119.1 | 215.6 | 279.5 |
| aHER2 N31A | 5.4E+04 | 6.5E-04 | 12 nM | 1.1E+03 | 95.8 | 265.8 | 485.8 |
| aHER2 S32A | 3.5E+04 | 5.4E-04 | 15 nM | 1.3E+03 | 231.7 | 222.9 | 402.2 |
| aHER2 C33A | N/A | N/A | N/A | N/A | N/A | 16.7 | 440.5 |
| aHER2 G34A | 1.3E+04 | 7.8E-04 | 59 nM | 8.9E+02 | 0.0 | 54.5 | 350.7 |
| aHER2 I52A | N/A | N/A | N/A | N/A | N/A | 30.3 | 404.3 |
| aHER2 S53A | 1.8E+04 | 2.2E-04 | 12 nM | 3.2E+03 | 481.4 | 233.4 | 298.9 |
| aHER2 G54A | 4.3E+04 | 2.4E-04 | 5.5 nM | 2.9E+03 | 25.8 | 105.7 | 205.1 |
| aHER2 D55A | 1.6E+04 | 3.5E-04 | 22 nM | 2.0E+03 | 450.3 | 210.9 | 339.3 |
| aHER2 G56A | 1.8E+04 | 2.4E-04 | 13 nM | 2.9E+03 | 249.9 | 150.7 | 337.9 |
| aHER2 D57A | 2.0E+04 | 3.6E-04 | 19 nM | 1.9E+03 | 469.9 | 267.5 | 365.7 |
| aHER2 T58A | 2.1E+04 | 2.9E-04 | 14 nM | 2.4E+03 | 373.9 | 244.0 | 294.1 |
| aHER2 V98A | 3.8E+04 | 5.9E-04 | 16 nM | 1.2E+03 | 156.9 | 156.3 | 187.3 |
| aHER2 C99A | N/A | N/A | N/A | N/A | N/A | 17.3 | 454.1 |
| aHER2 Y100A | 4.5E+04 | 3.4E-04 | 7.4 nM | 2.1E+03 | 130.6 | 163.9 | 271.9 |
| aHER2 N101A | 1.6E+04 | 2.8E-04 | 18 nM | 2.5E+03 | 443.3 | 226.6 | 400.9 |
| aHER2 L102A | 3.6E+04 | 1.4E-04 | 3.7 nM | 5.1E+03 | 49.9 | 101.9 | 176.5 |
| aHER2 E103A | 1.3E+04 | 4.3E-04 | 32 nM | 1.6E+03 | 450.1 | 198.5 | 443.5 |
| aHER2 T104A | 3.9E+03 | 2.5E-04 | 64 nM | 2.8E+03 | 219.1 | 65.4 | 514.3 |
| aHER2 Y105A | 3.0E+04 | 2.5E-04 | 8.4 nM | 2.8E+03 | 84.7 | 80.6 | 284.1 |
| aHER2 G27D | 2.0E+04 | 3.5E-04 | 17 nM | 2.0E+03 | 626.9 | 266.5 | 290.6 |
| aHER2 Y28D | 3.3E+04 | 2.7E-04 | 8.0 nM | 2.6E+03 | 308.6 | 203.7 | 168.6 |
| aHER2 I29D | 7.1E+04 | 2.0E-04 | 2.8 nM | 3.5E+03 | 17.4 | 180.8 | 343.6 |
| aHER2 F30D | 8.1E+04 | 5.6E-04 | 6.9 nM | 1.2E+03 | 44.5 | 207.2 | 270.8 |
| aHER2 N31D | 5.2E+04 | 6.7E-04 | 13 nM | 1.0E+03 | 147.8 | 221.4 | 259.4 |
| aHER2 S32D | 5.4E+04 | 6.6E-04 | 12 nM | 1.1E+03 | 69.2 | 170.8 | 260.2 |
| aHER2 C33D | N/A | N/A | N/A | N/A | N/A | 15.2 | 336.7 |
| aHER2 G34D | N/A | N/A | N/A | N/A | N/A | -5.4 | 265.6 |
| aHER2 I52D | N/A | N/A | N/A | N/A | N/A | 11.9 | 365.0 |
| aHER2 S53D | 2.6E+04 | 4.3E-04 | 17 nM | 1.6E+03 | 156.4 | 151.9 | 158.0 |
| aHER2 G54D | 8.4E+03 | 1.3E-04 | 16 nM | 5.3E+03 | 0.0 | 93.0 | 232.4 |
| aHER2 G56D | 2.0E+04 | 1.8E-04 | 8.8 nM | 3.9E+03 | 487.6 | 219.2 | 336.0 |
| aHER2 T58D | 2.1E+04 | 2.8E-04 | 13 nM | 2.5E+03 | 412.4 | 216.4 | 288.1 |
| aHER2 A97D | N/A | N/A | N/A | N/A | N/A | 6.2 | 390.0 |
| aHER2 V98D | N/A | N/A | N/A | N/A | N/A | 32.8 | 480.8 |
| aHER2 C99D | N/A | N/A | N/A | N/A | N/A | 12.6 | 371.4 |
| aHER2 Y100D | N/A | N/A | N/A | N/A | N/A | 168.0 | 222.7 |
| aHER2 N101D | 2.7E+04 | 2.5E-04 | 9.2 nM | 2.8E+03 | 651.5 | 220.4 | 210.5 |
| aHER2 L102D | N/A | N/A | N/A | N/A | N/A | 16.7 | 127.9 |
| aHER2 E103D | 2.4E+04 | 2.9E-04 | 12 nM | 2.4E+03 | 324.1 | 177.1 | 188.8 |
| aHER2 T104D | 9.1E+03 | 7.3E-05 | 8.1 nM | 9.4E+03 | 34.6 | 35.8 | 158.4 |

| | | | | | | | |
|---|---|---|---|---|---|---|---|
| aHER2 Y105D | N/A | N/A | N/A | N/A | N/A | 29.6 | 311.6 |
| aHER2 G27K | 2.1E+04 | 4.1E-04 | 20 nM | 1.7E+03 | 381.3 | 277.4 | 351.9 |
| aHER2 Y28K | 2.2E+04 | 2.5E-04 | 11 nM | 2.8E+03 | 169.4 | 125.4 | 128.6 |
| aHER2 I29K | 4.0E+04 | 5.1E-04 | 13 nM | 1.4E+03 | 178.8 | 216.9 | 473.9 |
| aHER2 F30K | 2.8E+04 | 4.8E-04 | 17 nM | 1.5E+03 | 484.7 | 333.2 | 396.5 |
| aHER2 N31K | 5.0E+04 | 5.5E-04 | 11 nM | 1.3E+03 | 66.2 | 136.3 | 249.6 |
| aHER2 S32K | 1.7E+04 | 4.8E-04 | 29 nM | 1.5E+03 | 205.1 | 184.4 | 369.1 |
| aHER2 C33K | N/A | N/A | N/A | N/A | N/A | 18.3 | 501.9 |
| aHER2 G34K | N/A | N/A | N/A | N/A | N/A | 5.8 | 455.0 |
| aHER2 I52K | 2.3E+04 | 4.1E-04 | 18 nM | 1.7E+03 | 104.7 | 70.6 | 478.3 |
| aHER2 S53K | N/A | N/A | N/A | N/A | N/A | 144.2 | 405.2 |
| aHER2 G54K | 7.3E+03 | 9.3E-05 | 13 nM | 7.5E+03 | 117.8 | 105.5 | 290.3 |
| aHER2 D55K | 2.1E+04 | 4.6E-04 | 22 nM | 1.5E+03 | 136.6 | 175.8 | 401.5 |
| aHER2 G56K | 2.6E+04 | 4.9E-04 | 19 nM | 1.4E+03 | 182.1 | 166.0 | 505.2 |
| aHER2 D57K | 1.2E+04 | 4.4E-04 | 35 nM | 1.6E+03 | 195.8 | 158.7 | 235.2 |
| aHER2 T58K | 1.6E+04 | 2.1E-04 | 13 nM | 3.3E+03 | 391.5 | 196.7 | 239.2 |
| aHER2 A97K | N/A | N/A | N/A | N/A | N/A | 25.2 | 280.5 |
| aHER2 V98K | 3.8E+04 | 5.1E-04 | 13 nM | 1.4E+03 | 50.1 | 162.1 | 370.2 |
| aHER2 C99K | N/A | N/A | N/A | N/A | N/A | 29.8 | 366.9 |
| aHER2 Y100K | N/A | N/A | N/A | N/A | N/A | 98.0 | 251.4 |
| aHER2 N101K | 1.5E+04 | 2.4E-04 | 16 nM | 2.9E+03 | 329.3 | 232.7 | 380.6 |
| aHER2 L102K | N/A | N/A | N/A | N/A | N/A | 89.5 | 249.2 |
| aHER2 E103K | 1.6E+04 | 2.0E-04 | 12 nM | 3.5E+03 | 89.7 | 92.4 | 371.0 |
| aHER2 T104K | N/A | N/A | N/A | N/A | N/A | -7.3 | 361.6 |
| aHER2 Y105K | 2.0E+04 | 4.4E-04 | 21 nM | 1.6E+03 | 98.2 | 98.7 | 306.1 |
| aHER2 G27S | 2.0E+04 | 3.6E-04 | 18 nM | 1.9E+03 | 666.6 | 313.4 | 393.1 |
| aHER2 Y28S | 1.8E+04 | 2.6E-04 | 14 nM | 2.7E+03 | 559.1 | 208.4 | 237.6 |
| aHER2 I29S | N/A | N/A | N/A | N/A | N/A | 44.9 | 36.4 |
| aHER2 F30S | 2.1E+04 | 6.2E-04 | 30 nM | 1.1E+03 | 192.4 | 57.8 | 54.6 |
| aHER2 N31S | 3.9E+04 | 4.8E-04 | 12 nM | 1.4E+03 | 50.2 | 95.6 | 130.9 |
| aHER2 C33S | N/A | N/A | N/A | N/A | N/A | 18.2 | 338.0 |
| aHER2 G34S | N/A | N/A | N/A | N/A | N/A | 49.5 | 281.2 |
| aHER2 I52S | N/A | N/A | N/A | N/A | N/A | 27.2 | 419.6 |
| aHER2 S53S | 2.0E+04 | 2.7E-04 | 13 nM | 2.6E+03 | 317.7 | 254.6 | 304.8 |
| aHER2 G54S | 1.5E+04 | 7.1E-05 | 4.8 nM | 9.8E+03 | 53.9 | 142.8 | 295.4 |
| aHER2 D55S | 2.0E+04 | 3.4E-04 | 18 nM | 2.0E+03 | 222.8 | 267.1 | 406.6 |
| aHER2 G56S | 1.7E+04 | 3.3E-04 | 19 nM | 2.1E+03 | 210.7 | 210.9 | 399.9 |
| aHER2 D57S | 2.5E+04 | 4.8E-04 | 19 nM | 1.5E+03 | 218.3 | 226.1 | 314.6 |
| aHER2 T58S | 1.9E+04 | 3.3E-04 | 18 nM | 2.1E+03 | 413.2 | 218.0 | 256.0 |
| aHER2 A97S | 9.4E+04 | 4.2E-04 | 4.5 nM | 1.6E+03 | 36.0 | 180.0 | 263.1 |
| aHER2 V98S | 4.1E+04 | 5.6E-04 | 14 nM | 1.2E+03 | 20.8 | 138.8 | 353.3 |
| aHER2 C99S | N/A | N/A | N/A | N/A | N/A | 21.2 | 359.0 |
| aHER2 Y100S | 5.6E+04 | 4.2E-04 | 7.5 nM | 1.6E+03 | 100.3 | 187.9 | 221.8 |
| aHER2 N101S | 2.0E+04 | 3.5E-04 | 18 nM | 2.0E+03 | 338.8 | 230.3 | 383.6 |
| aHER2 L102S | 5.5E+04 | 2.3E-04 | 4.1 nM | 3.0E+03 | 10.5 | 115.7 | 169.5 |
| aHER2 E103S | 1.1E+04 | 3.8E-04 | 34 nM | 1.8E+03 | 360.9 | 162.7 | 326.1 |
| aHER2 T104S | 3.6E+04 | 3.0E-04 | 8.2 nM | 2.3E+03 | 23.5 | 84.3 | 201.4 |
| aHER2 Y105S | 2.6E+04 | 5.1E-04 | 20 nM | 1.4E+03 | 165.4 | 116.3 | 260.8 |
